# Deficiency of the *ywhaz* gene, involved in neurodevelopmental disorders, alters brain activity and behaviour in zebrafish

**DOI:** 10.1101/2021.06.30.450513

**Authors:** Ester Antón-Galindo, Elisa Dalla Vecchia, Javier G Orlandi, Gustavo Castro, Emilio J Gualda, Andrew MJ Young, Fernando Aguado, Pablo Loza-Alvarez, Bru Cormand, William HJ Norton, Noèlia Fernàndez-Castillo

## Abstract

Genetic variants in *YWHAZ* contribute to psychiatric disorders such as autism spectrum disorder and schizophrenia, and have been related to an impaired neurodevelopment in humans and mice. Here, we used zebrafish to further understand the mechanisms by which *YWHAZ* contributes to neurodevelopmental disorders. We observed that *ywhaz* expression was panneuronal during developmental stages and restricted to Purkinje cells in the adult cerebellum, cells that are described to be reduced in number in autistic patients. We then performed whole-brain imaging in wild-type and *ywhaz* CRISPR/Cas9 knockout (KO) larvae and found altered neuronal activity and connectivity in the hindbrain. Adult *ywhaz* KO fish display decreased levels of monoamines in the hindbrain and freeze when exposed to novel stimuli, a phenotype that can be reversed with drugs that target monoamine neurotransmission. These findings suggest an important role for *ywhaz* in establishing neuronal connectivity during development and modulating both neurotransmission and behaviour in adults.

## INTRODUCTION

*YWHAZ* is a member of the 14-3-3 gene family, coding for a highly conserved group of molecular chaperones that play important roles in biological processes and neuronal development (Cornell and Toyo-oka, 2017). *YWHAZ* encodes 14-3-3ζ, a protein involved in neurogenesis and neuronal migration as shown by morphological changes in the brain of 14-3-3ζ knockout mice (Cheah et al., 2012; Jaehne et al., 2015; Toyo-Oka et al., 2014; Xu et al., 2015). These animals present behavioural and cognitive alterations that have been related to psychiatric disorders like schizophrenia (Cheah et al., 2012; Xu et al., 2015), including hyperactivity, impaired memory, lower anxiety and impaired sensorimotor gating.

In humans, several studies have pointed to an association between Y*WHAZ* and psychiatric disorders. Genetic studies have associated *YWHAZ* polymorphisms to major depression and schizophrenia (Jia et al., 2004; Torrico et al., 2020). In addition, a protein-protein interaction analysis including all genes found mutated in an exome sequencing study of ASD multiplex families reported YWHAZ as the main node of the network (Toma et al., 2014) and a heterozygous frameshift mutation in the *YWHAZ* gene, inherited from a mother with depression, was found to have functional implications in two siblings diagnosed with autism spectrum disorder (ASD) and attention deficit/hyperactivity disorder (ADHD) (Torrico et al., 2020). Furthermore, decreased expression of the *YWHAZ* gene and low levels of the corresponding 14-3-3ζ protein were reported in *postmortem* brains of patients with neurodevelopmental disorders like ASD and schizophrenia (English et al., 2011; Middleton et al., 2005; Torrico et al., 2020), and 14-3-3 protein levels are reduced in platelets and pineal glands of ASD patients (Pagan et al., 2014, 2017). However, even though evidence points to a contribution of *YWHAZ* to neurodevelopmental and psychiatric disorders, the mechanisms underlying its contribution remain unclear and further studies need to be performed to understand the role of *YWHAZ* in brain development and function.

A homologue of *YWHAZ* is also found in zebrafish, a species that represents a powerful model to study brain development and psychiatric disorders (Kalueff et al., 2014; Norton, 2013; Vaz et al., 2019). Zebrafish display well-defined behaviours that can be translated to humans in some cases (Kalueff et al., 2014; Vaz et al., 2019) and, although neurodevelopmental timings and brain organization differ between human and zebrafish, comparative studies have precisely mapped these differences and the information gained in zebrafish can be transferred to other species (Kalueff et al., 2014; Kozol et al., 2016). Importantly, zebrafish present a high genetic homology to humans, are easy to manipulate genetically and their small size and transparency during larval stages make zebrafish ideal for *in vivo* imaging studies (Kalueff et al., 2014; Norton, 2013). Indeed, whole-brain imaging is a recently developed technique (Ahrens et al., 2013; Vanwalleghem et al., 2018) that allows *in vivo* investigation of neuronal activity and connectivity in zebrafish using light-sheet microscopy (Olarte et al., 2018). This novel approach combined with genetic engineering represents an excellent tool to use in specific zebrafish models to study the neural mechanisms of human brain disorders.

In this study, we aimed to investigate the role of *YWHAZ* in brain development and function using a novel mutant line. We explored the effect of loss of *ywhaz* function on neural activity and connectivity during development, and in neurotransmission and behaviour during adulthood.

## RESULTS

### *ywhaz* expression is pan-neuronal during development and restricted to the cerebellum in adults

We first investigated *ywhaz* gene expression in the developing brain using *in situ* hybridization (ISH). In three to nine days post-fertilization (dpf) WT larvae, *ywhaz* expression is widespread covering almost all brain areas, with a particularly strong signal in the cerebellum of whole mount embryos (Figure 1A). In contrast, in adult zebrafish, *ywhaz* expression is restricted to the cerebellum. *ywhaz* RNA is present in the granule cell layer (GCL) of the valvula cerebelli (Va) and crista cerebellaris (CCe), in an area that appears to be the Purkinje cell layer (PCL) (Figure 1B).

**Figure 1.**
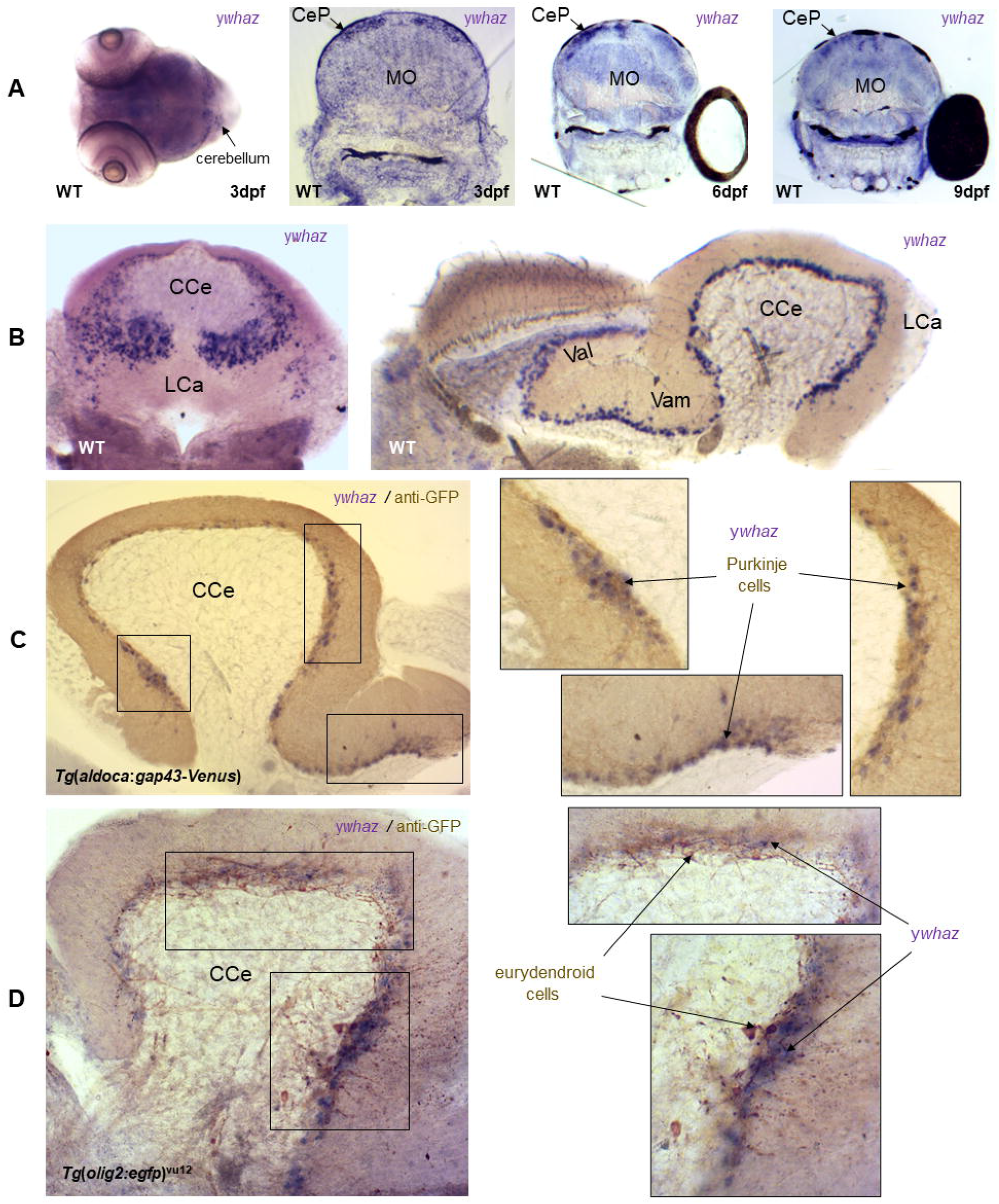
*ywhaz* mRNA is widely expressed during zebrafish development but restricted to the Purkinje cells in the cerebellum in adults. (A) *In situ* hybridization (ISH) using a *ywhaz* RNA probe (purple) of 3-9 days post-fertilization (dpf) WT zebrafish embryos shows that *ywhaz* expression is widespread covering almost all brain areas, with a strongest signal in cerebellum. On the left, a whole-mount of a 3 dpf embryo. On the right, coronal sections of 3, 6 and 9 dpf embryos, including the MO and CeP brain areas. (B) Coronal and sagittal sections of WT adult brains after ISH with a *ywhaz* RNA probe (purple) shows that in WT adult brains, *ywhaz* expression is restricted to the granule cell layer in the cerebellum. (C) Co-staining of the adult *Tg(aldoca:gap43-Venus)* cerebellum with a *ywhaz* riboprobe (purple) by ISH and anti-GFP antibody (brown) by immunohistochemistry (IHC) on a sagittal section. The overlap between the two staining (black arrows) indicates that *ywhaz* is localised within Purkinje cells. (D) Co-staining of the adult *Tg*(*olig2*:*egfp*)^vu12^ cerebellum with a *ywhaz* riboprobe (purple) by ISH and anti-GFP antibody (brown) by IHC on a sagittal section. The absence of overlap between the two stains shows that *ywhaz* mRNA is not localized in eurydendroid cells. CCe, corpus cerebelli; CeP, cerebellar plate; LCa, lobus caudalis cerebelli; MO, medulla oblongata; Val, lateral valvula cerebelli; Vam, medial valvula cerebelli.

We next combined *ywhaz* ISH with an anti-GFP antibody stain on adult brain sections of *Tg*(*aldoca:gap43-Venus*) (Ahn et al., 1994) and *Tg*(*olig2*:*egfp*)^vu12^ (McFarland et al., 2008) transgenic lines, to confirm that *ywhaz* is expressed only within Purkinje cells and not in eurydendroid cells, the zebrafish equivalent of the deep cerebellar nuclei in humans (Hibi and Shimizu, 2012). In the *Tg*(*aldoca:gap43-Venus*) line, the promoter of the *aldolase Ca* (*aldoca*) gene (Ahn et al., 1994) is used as a driver and EGFP labels Purkinje cells, inhibitory neurons in the PCL. In the *Tg*(*olig2*:*egfp*)^vu12^ fish cerebellum, EGFP labels eurydendroid cells (McFarland et al., 2008), excitatory neurons that are situated ventrally to Purkinje cells (Biechl et al., 2016). We found that *ywhaz* ISH staining overlaps GFP in the *Tg*(*aldoca:gap43-Venus*) line, meaning that *ywhaz* is expressed within Purkinje cells and not within eurydendroid cells at adult stages (Figure 1C and D).

### Generation of a *ywhaz*^*-/-*^ zebrafish mutant line by CRISPR-Cas9 mutagenesis

To assess the role of *ywhaz* during neural development we generated a novel *ywhaz* mutant line (Kai et al., 2021). We used CRISPR/Cas9 to engineer a mutation within the third exon of *ywhaz*. We selected a F0 fish carrying a 7-bp deletion (380_387delCCTGGCA) as a founder to generate a stable *ywhaz* mutant line (Figure S1). This deletion causes a frameshift that leads to a premature stop codon in the third exon of the unique *ywhaz* isoform, generating an ORF that is 48% shorter than the normal one (Figure S1C and D). The F1 embryos born from the cross between the F0 founder and WT were raised to adulthood, genotyped, and only the F1 fish carrying the 7-bp deletion were kept.

To check whether the 7-bp deletion reduces *ywhaz* expression we performed ISH in larvae and adult brains and confirmed loss of *ywhaz* expression at the mRNA level in the brain of *ywhaz*^*-/-*^ fish, at both developmental (Figure 2A-C) and adult stages (Figure 2D). In addition, to confirm that the deletion reduces mRNA levels in zebrafish brains we analysed *ywhaz* mRNA levels by RT-qPCR. We observed a significantly decreased level of *ywhaz* expression in *ywhaz*^-/-^ compared to WT (p < 0.0001, Mann-Whitney U test, Figure 2E), which suggests that nonsense-mediated mRNA decay degradation (Karousis et al., 2016) of the truncated *ywhaz* transcript has occurred in mutants. We also analysed *ywhae* mRNA levels, another member of the 14-3-3 gene family, and observed a difference in expression of *ywhae* in *ywhaz*^-/-^ brains that does not reach significance compared to WT (Medians: WT = 2.18, *ywhaz*^-/-^ = 1.062, p = 0.075, Mann-Whitney U test, Figure 2F), probably due to compensatory mechanisms.

**Figure 2.**
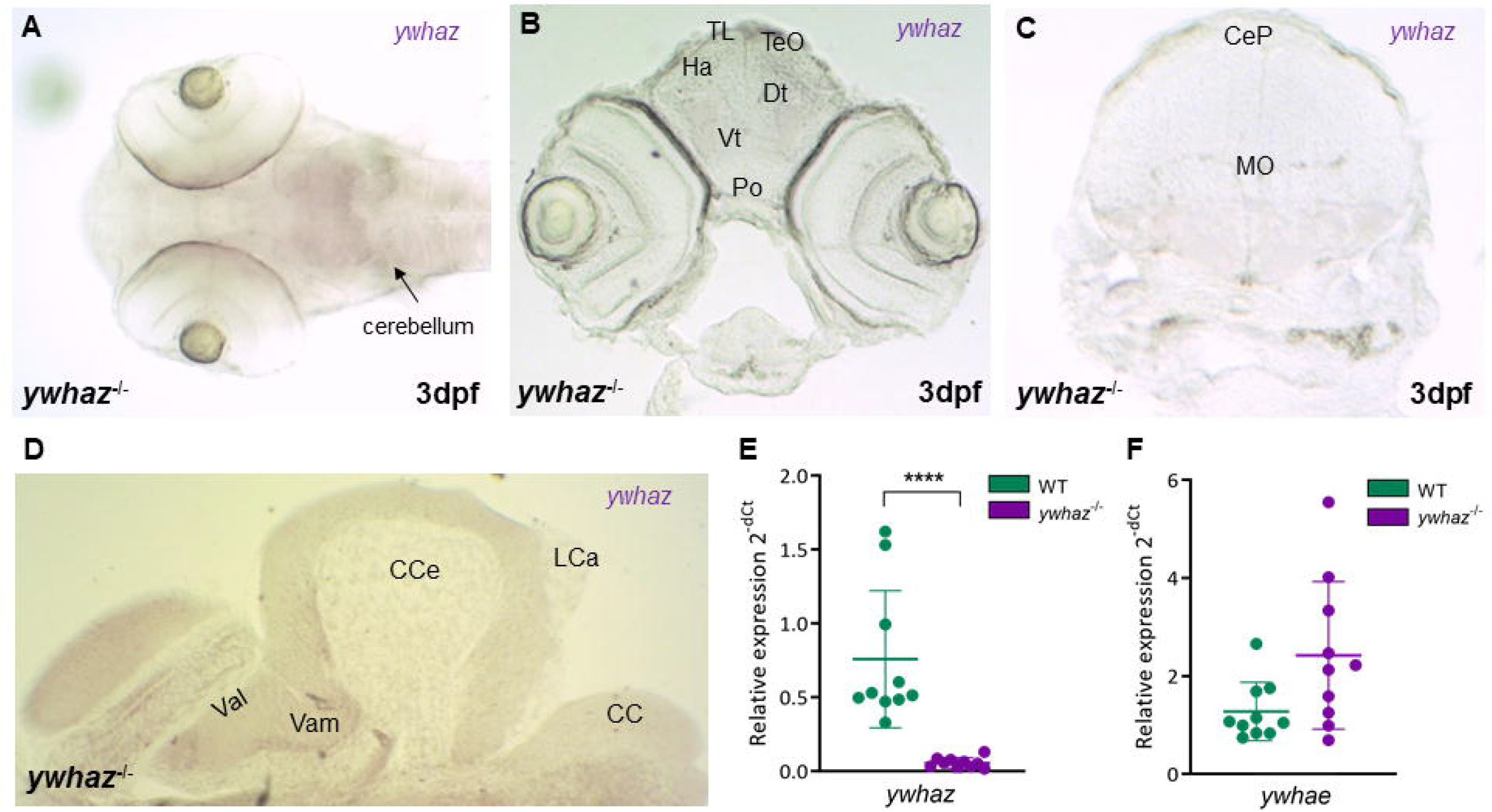
*ywhaz* is not expressed in *ywhaz*^-/-^ embryos and *ywhaz*^-/-^ adult brains. (A-C) *In situ* hybridization of 3 days post-fertilization (dpf) larvae reveals that *ywhaz* is not expressed during development in ywhaz^-/-^. (D) *In situ* hybridization of adult brains reveals that *ywhaz* is not expressed during adulthood in *ywhaz*^-/-^. (E) RT-qPCR amplification demonstrates that *ywhaz* expression is marginal in the brain of *ywhaz*^-/-^ adults compared to WT. Relative expression profile of *ywhaz* normalised to the reference gene elongation factor 1a (*elf1a*). p < 0.0001, Mann-Whitney U test, n=10 WT, n=10 ywhaz^-/-^. (F) RT-qPCR amplification shows a trend increase in *ywhae* expression in the brain of *ywhaz*^-/-^ adults compared to WT although the difference is not significant. Relative expression profile of *ywhae* normalised to the reference gene elongation factor 1a (*elf1a*). p = 0.075, Mann-Whitney U test, n=10 WT, n=10 ywhaz^-/-^. **** p<0.0001. Mean ± SD. CC, crista cerebellaris; CCe, corpus cerebelli; CeP, cerebellar plate; Dt, dorsal thalamus; Ha, habenula; LCa, lobus caudalis cerebelli; MO, medulla oblongata; Po, preoptic region; TL, torus longitudinalis; TeO, optic tectum; Val, lateral valvula cerebelli; Vam, medial valvula cerebelli, Vt, ventral thalamus.

### *ywhaz* deficiency alters spontaneous neuronal activity and functional connectivity in the hindbrain of larvae

The pan-neuronal expression pattern of *ywhaz* suggests that it may play a role during neural development. We performed whole-brain imaging *in vivo* at 6 dpf to investigate changes in neural circuit function and connectivity (see Figure 3 for methodological details and Supplementary Videos 1-3).

**Figure 3.**
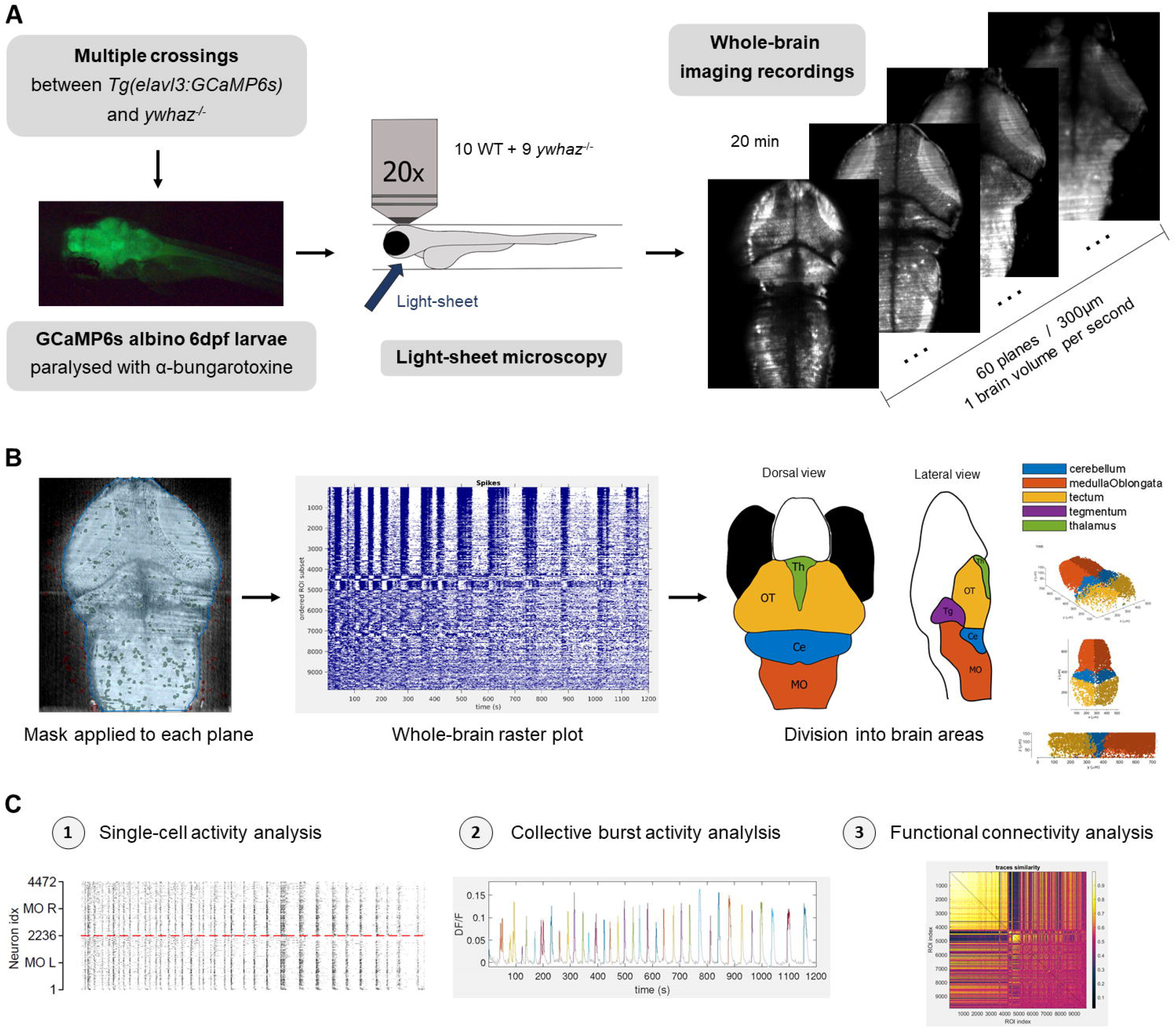
Whole-brain imaging methods. (A) Steps followed to perform the whole-brain imaging recordings in 6 days post-fertilization (dpf) larvae expressing GCaMP6s pan-neuronally. First, multiple crossings were performed between the *Tg*(*elavl3:GCaMP6s*) and *ywhaz*^-/-^ mutant lines in order to obtain *ywhaz*-deficient albino larvae expressing GCaMP6s pan-neuronally. Then, *Tg*(*elavl3:GCaMP6s*) *ywhaz*^-/-^ and *Tg*(*elavl3:GCaMP6s*) *ywhaz*^+/+^ 6 dpf larvae were paralysed with α-bungarotoxine and placed in a plastic tube to perform the whole-brain imaging recordings with a light-sheet microscope. (B) Steps followed in the single-cell fluorescence traces extraction. First, a mask was applied to each single plane to avoid detection of fluorescence outside the brain. Then, single-cell fluorescence traces were extracted for all the neurons detected in each single plane and combined to obtain neuronal activity from the whole-brain of each individual during the 20 min recording. Finally, brain regions were manually defined in order to analyse them separately. On the right, plots of all the neurons detected in each brain area of one individual. In a first approach, we divided the three largest regions (Ce, OT, and MO) by hemispheres and obtained no differences in the number of neurons or activity between hemispheres. Ce, cerebellum; MO, medulla oblongata; OT, optic tectum; ROI, region of interest; Te, tegmentum; Th, thalamus. (C) Three levels of analysis were performed in each of the five defined brain areas: single-cell activity, collective burst activity, and functional connectivity.

We identified differences in hindbrain spontaneous activity and functional connectivity between the WT and *ywhaz*^-/-^ genotypes. We found an increased number of active neurons in medulla oblongata (MO) of *ywhaz*^*-/-*^ fish (p = 0.046, Figure 4A), which represents a higher fraction of total active neurons (p = 0.005, Figure 4B). This relative increase of active neurons in the MO correlates with a significant decrease in active cerebellar neurons in *ywhaz*^-/-^ fish (p = 0.018, Figure 4B). In thalamus and tegmentum, the low number of active neurons that we could detect prevented us from properly assessing activity and connectivity in these areas (Figure 4A and B).

**Figure 4.**
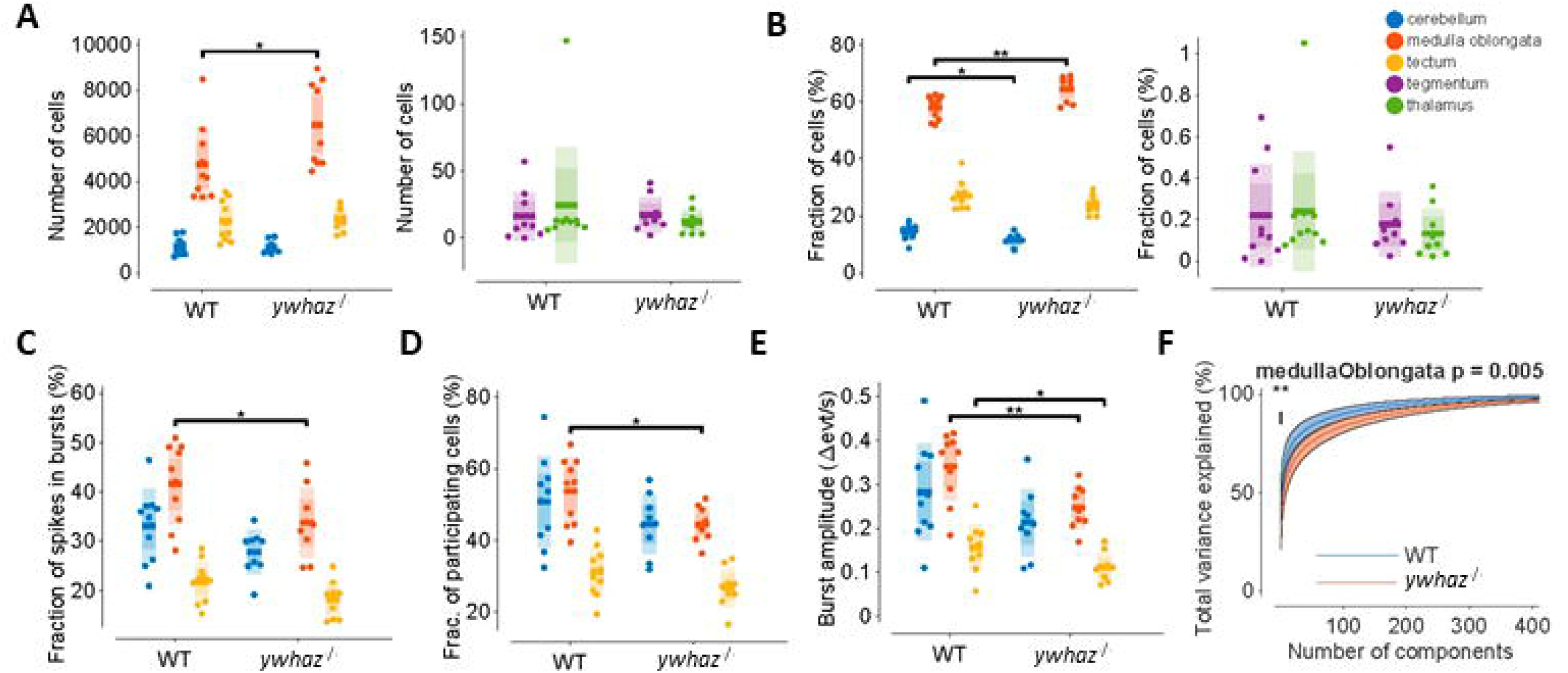
Decreased coherence in the hindbrain neuronal activity in zebrafish *ywhaz*^-/-^ larvae. (A) Number of active neurons detected in each brain area. (B) Fraction of total active neurons detected in each brain area. (E) Fraction of total spikes participating in single-cell bursts in each brain area. (F) Fraction of total cells participating in collective bursts in each brain area. (G) Burst amplitude in each brain area, measured as the increase of spikes (events) per second compared to basal spike activity. For A-G datasets: unpaired t-tests, each single point represents an individual, the central line represents the mean, the coloured darker shadow represents the 95% confidence interval and the lighter shadow the standard deviation. (H) Principal component analysis of the neuronal activity in the medulla oblongata (MO). The central line represents the mean and the coloured shadow represents the 95% confidence interval. * p < 0.05, ** p < 0.01. n= 10 WT and 9 *ywhaz*^*-/-*^.

Single-cell activity analysis (Figure S2) showed that a lower fraction of total spikes was contained within single-cell bursts in the MO of *ywhaz*^*-/-*^ fish (p = 0.039, Figure 4C). In addition, population-level analysis (Figure S3) showed that a lower fraction of neurons participated in large-scale bursts (defined as large increases in activity within an area) in the MO of *ywhaz*^*-/-*^ fish (p = 0.016, Figure 4D), and the average increase in activity during these bursts was also lower in MO and tectum in *ywhaz*^*-/-*^ fish (p = 0.006 for MO and p = 0.043 for tectum, Figure 4E). Finally, principal component analysis (PCA) showed that for any target variance explained, *ywhaz*^-/-^ always required a larger number of components, suggesting less coherence in the activity modes of *ywhaz*^-/-^ in the MO (p = 0.005, Figure 4F). This effect was not found in other areas (Figure S4A). We then explored possible alterations in functional connectivity inside each of the defined areas (Figure S4B). We found a higher clustering coefficient (relative number of loops in a network) (p = 0.022, Figure 5A) and a higher global efficiency (a measure of how easy it is for activity to move throughout a network) (p = 0.044, Figure 5B) in cerebellum of *ywhaz*^*-/-*^ fish. Also, we observed a higher assortativity (the correlation between the number of connections of connected neurons) (p = 0.001, Figure 5C) and a higher Louvain community statistic (a measure of how segregated the communities of a network are) in MO of *ywhaz*^*-/-*^fish (p = 0.010, Figure 5D). Finally, analyses of connectivity distribution showed that in the MO of WT fish there is a subpopulation of highly connected neurons (connected with 30-40% of the MO neurons) that is not present in the MO of *ywhaz*^*-/-*^fish, but no differences were found in cerebellum (Figure 5E and F). Given the other measures, these highly connected neurons in the MO of WT fish are likely to be responsible for connecting communities within a network together, resulting in a more coherent pattern of collective activity in WT fish, and their absence in *ywhaz*^-/-^ fish would lead to an impairment of MO activity.

**Figure 5.**
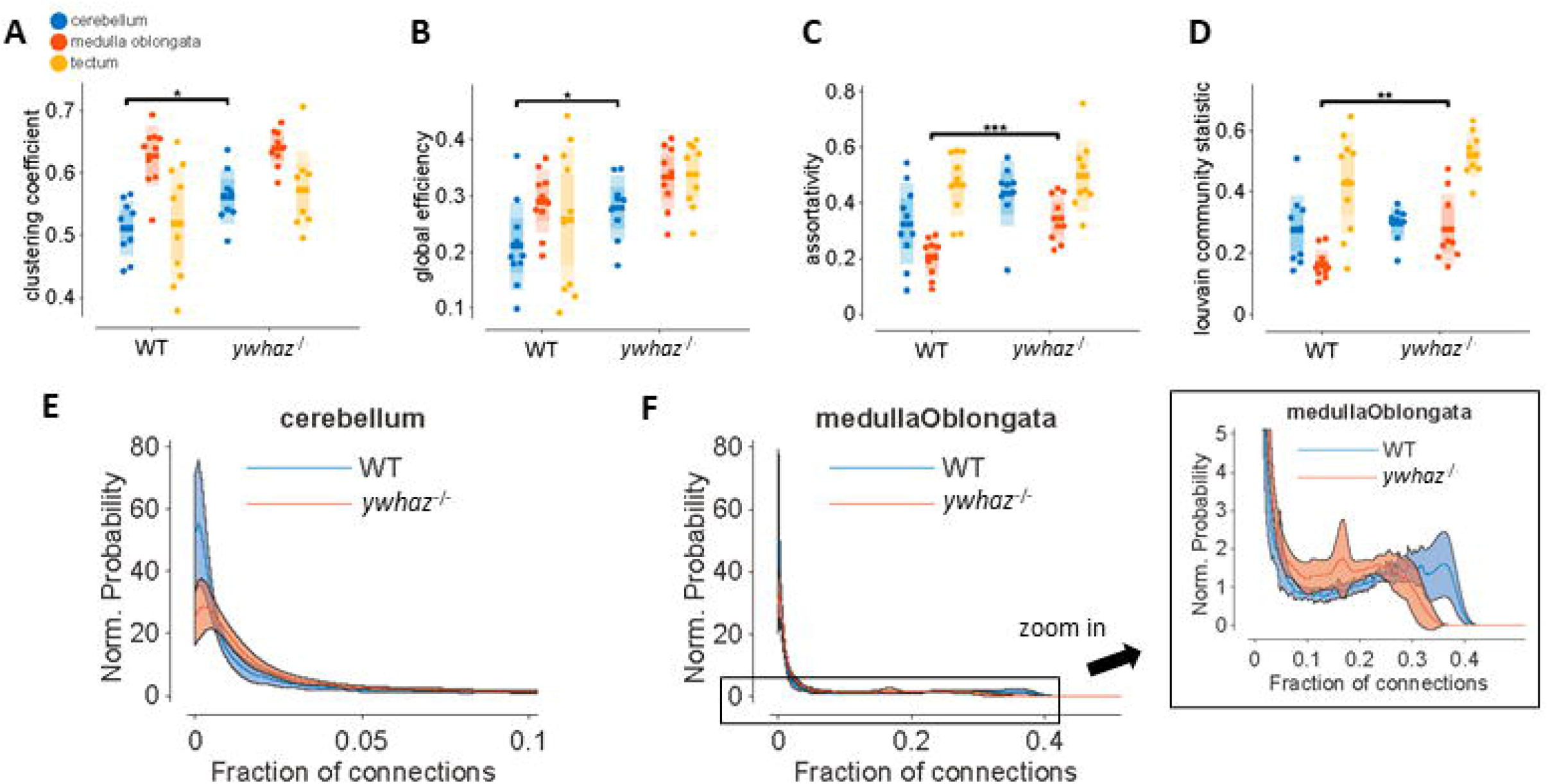
Decreased hindbrain neuronal connectivity in zebrafish *ywhaz*^-/-^ larvae. (A) Normalized clustering coefficient in each brain area. (B) Global efficiency in each brain area. (C) Assortativity in each brain area. (D) Louvain community statistic in each brain area. For A-D datasets: unpaired t-tests, each single point represents an individual, the central line represents the mean, the darker bar the 95% confidence interval and the lighter bar the standard deviation. (E) No significant differences were found in the connectivity distribution in the cerebellum of WT and *ywhaz*^-/-^ larvae. (F) Connectivity distribution in the medulla oblongata (MO) of WT and *ywhaz*^-/-^ larvae. A subpopulation of highly connected neurons exists in the MO of WT larvae that is absent in *ywhaz*^-/-^ larvae. For E and F: the central line represents the mean and the coloured shadow represents the 95% confidence interval. YWHAZ, *ywhaz*^-/-^ larvae; WT, wild-type larvae. n= 10 WT and 9 *ywhaz*^*-/-*^. * p < 0.05, ** p < 0.01, *** p < 0.001.

Altogether, these results show that in *ywhaz*^*-/-*^ fish cerebellar neurons represent a lower fraction of total brain neurons but present a higher clustering and more effective connectivity. Also, these results point to the presence of more isolated neuronal communities in the MO of *ywhaz*^*-/-*^ fish, which generates a lower collective burst activity and synchrony in this area and would lead to an impaired MO activity. Spontaneous synchronized neuronal communication plays an important role during early development in establishing the mature brain circuitry, not only in humans but also in other vertebrates including zebrafish (Avitan et al., 2017; Marachlian et al., 2018; Molnár et al., 2020; Momose-Sato and Sato, 2016). We therefore hypothesized that the alterations in this spontaneous activity may affect neuronal migration and wiring and have a long-term impact in neurotransmission.

### *ywhaz* deficiency alters monoamine levels in adult hindbrain

We next investigated whether the observed alterations in neuronal activity and connectivity caused by a loss of function of *ywhaz* gene during development might affect neurotransmitter signalling in adults. We performed high performance liquid chromatography (HPLC) to measure the basal levels of several neurotransmitters: 3,4-dihydroxyphenylacetic acid (DOPAC), dopamine (DA), 5-hydroxyindoleacetic acid (5HIAA) and serotonin (5-HT), in the brain of *ywhaz*^-/-^ and WT adult fish. We found a significant reduction of DA and 5-HT levels in the hindbrain of *ywhaz*^-/-^ (p = 0.0063 and p = 0.0026 respectively, Figure 6A). No further alterations were found in other areas of the brain (Figure S5A-C) nor in the breakdown of 5-HT to 5HIAA, or DA to DOPAC (Figure S5D-E).

**Figure 6.**
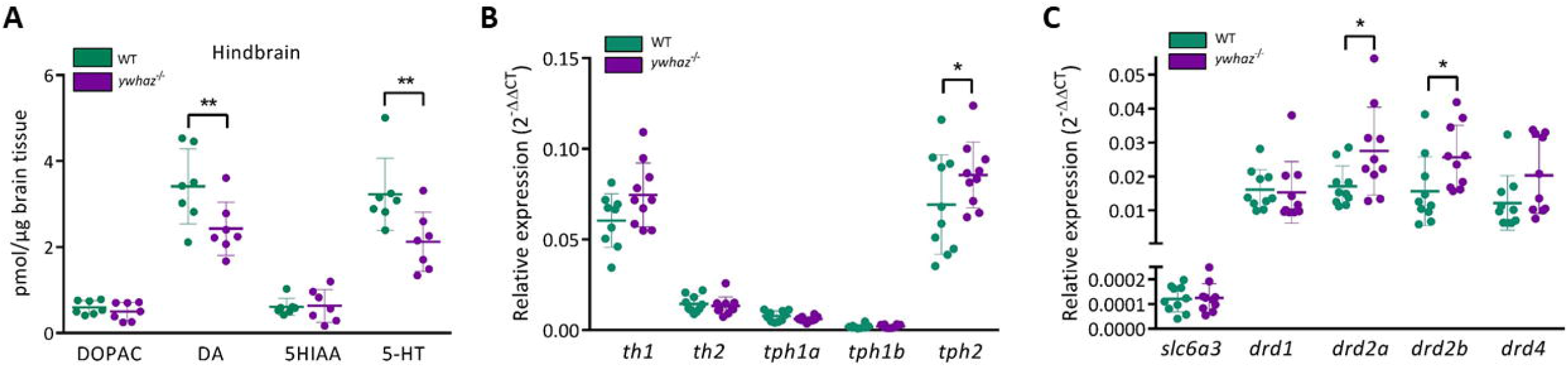
Alterations in the monoamine neurotransmission in the hindbrain of adult KO. (A) High performance liquid chromatography in the hindbrain of WT and *ywhaz*^-/-^ adult zebrafish. DA and 5-HT levels are decreased in the hindbrain of *ywhaz*^-/-^ compared to WT. DA, dopamine; DOPAC, 3,4-dihydroxyphenylacetic acid; 5-HIAA, 5-hydroxyindoleacetic acid; 5-HT, 5-hydroxytryptamine. n = 7 WT, n = 7 ywhaz^-/-^. (B) Relative expression profile of *tyrosine hydroxylase 1* (*th1*), *tyrosine hydroxylase 2* (*th2*), *tryptophan hydroxylase 1a* (*tph1a*), *tryptophan hydroxylase 1b* (*tph1b*) and *tryptophan hydroxylase* 2 (*tph2*), normalised to the reference gene ribosomal protein L13 (*rpl13*). *ywhaz*^-/-^ fish have an increased level of *tph2* expression. n = 10 WT, n = 10 *ywhaz*^-/-^. (C) Relative expression profile of the neurotransmitter transporter *solute carrier family 6 member 3* (*slc6a3*) and dopamine receptors *dopamine receptor 1* (*drd1*), *dopamine receptor 2a* (*drd2a*), *dopamine receptor 2b* (*drd2b*) and *dopamine receptor 4* (*drd4*), normalised to the reference gene *elongation factor 1a* (*elf1a*). *ywhaz*^-/-^ have an increased level of *drd2a* and *drd2b* expression. n = 10 WT, n = 10 *ywhaz*^-/-^. Multiple t-tests with Holm-Sidak correction for multiple comparisons.

### Loss of *ywhaz* function alters the expression of genes involved in the dopaminergic and serotonergic pathways

The reduced levels of DA and 5-HT in mutants suggests that *ywhaz* may influence the synthesis of these neurotransmitters. Tyrosine hydroxylase (TH) and Tryptophan hydroxylase (TPH), rate-limiting enzymes in the biosynthesis of DA and 5-HT, respectively, are known to be regulated by 14-3-3 proteins (Aitken, 2006). We therefore first quantified the expression of genes coding for the DA and 5-HT synthesis enzymes TH and TPH in adult fish. There was an increase in *tryptophan hydroxylase 2* (*tph2*) and *tyrosine* hydroxylase 1 (*th1*) expression in *ywhaz*^-/-^ compared to WT, although differences in *th1* did not overcome testing for multiple corrections. *tph1a, tph1b* and *th2* showed all a low expression in both genotypes (Figure 6B). We further examined the transcription of genes coding for proteins involved in the dopaminergic neurotransmitter pathway: the dopamine transporter *solute carrier family 6 member 3* (*slc6a3*), *dopamine receptor 1* (*drd1*), *dopamine receptor 2a* (*drd2a*), *dopamine receptor 2b* (*drd2b*) and *dopamine receptor 4* (*drd4*). The expression of the two isoforms *drd2a and drd2b* was significantly increased in *ywhaz*^-/-^ compared to WT (Figure 6C).

### Monoaminergic drugs reverse *ywhaz*^-/-^ freeze in response to novelty

We performed a battery of behavioural tests in adult WT and *ywhaz*^-/-^ mutants to characterize possible alterations in behaviour due to *ywhaz* deficiency (Figure S6). We only found differences between genotypes in the visually-mediated social preference (VMSP) test, used to assess social behaviour. During the social preference step, both genotypes spent most of the time swimming close to the first group of strangers (p < 0.0001 for both WT and *ywhaz*^-/-^; Figure S6D). However, during the preference for social novelty step, WT switched preference to the second group of unfamiliar fish (p = 0.0475, Figure S6D) whereas *ywhaz*^-/-^ fish did not show preference between the two groups of strangers (p = 0.98, Figure S6D). Instead, we found that immediately after the addition of the second group of unfamiliar fish, *ywhaz*^-/-^ mutants froze significantly more than WT (p = 0.0035; Figure 7A). To test whether it was the live stimulus that caused this phenotype, we repeated the test by adding marbles instead of the second group of unfamiliar fish and observed the same freezing behaviour in *ywhaz*^-/-^ mutants (p = 0.0251, Figure 7B). Contrary to adults, juvenile *ywhaz*^-/-^ fish did not show an altered social behaviour: they did not show any increase in freezing behaviour in the VMSP test (p > 0.99, Figure S6F), nor an altered cluster score in the shoaling test for social interaction in a group (Figure S6G).

**Figure 7.**
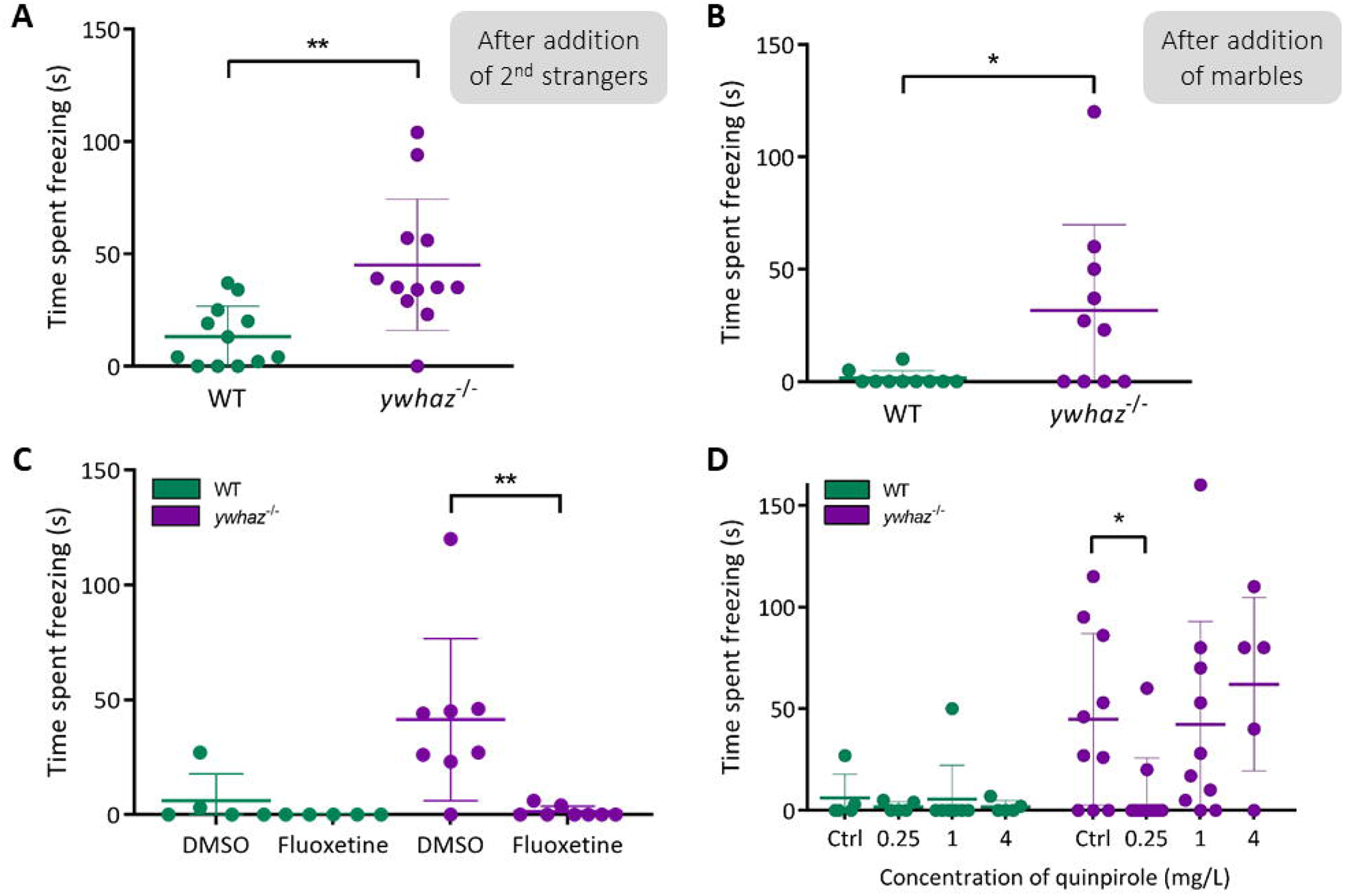
Treatment with fluoxetine and quinpirole reverses the freezing behaviour observed in *ywhaz*^-/-^ mutants. (A and B) Time spent freezing during the first two minutes after the addition of the second group of strangers (A) or marbles (B) in the behavioural setup during the second step of the visually mediated social preference test. Strangers: Unpaired t-test with Welch’s correction; n = 12. Marbles: Mann-Whitney U test; n = 10. (C and D) Treatment with 5 mg/L fluoxetine and 0.25 mg/L quinpirole rescues the freezing phenotype in *ywhaz*^-/-^. (C) Time spent freezing after the addition of the second group of unfamiliar fish in the tank with WT or *ywhaz*^-/-^ fish treated with 5 mg/L fluoxetine or DMSO. Multiple t-tests with Holm-Sidak correction for multiple comparisons. n = 5 WT, n = 8 *ywhaz*^-/-^. (D) Time spent freezing after the addition of the second group of unfamiliar fish in the tank with WT or *ywhaz*^-/-^ fish treated with different concentrations of quinpirole. Kruskal-Wallis test with Dunn’s multiple comparisons. n = 5 WT and n = 10 *ywhaz*^-/-^ for Ctrl, 0.25 mg/L and 1 mg/L; and n = 5 *ywhaz*^-/-^ for 4 mg/L. ** p < 0.01, * p < 0.05. Mean ± standard deviation.

HPLC and qPCR results suggested that alterations in 5-HT and DA signalling may underlie the behavioural phenotype of *ywhaz*^-/-^. We therefore used fluoxetine, a selective serotonin reuptake inhibitor (SSRI) (Vaswani et al., 2003), and quinpirole, a selective D2-like receptor agonist (Millan et al., 2002) to investigate the connection between 5-HT, DA, and the behavioural phenotype observed in *ywhaz*^-/-^ in the VMSP test. Treatment with 5 mg/L fluoxetine and treatment with 0.25 mg/L quinpirole significantly decreased the time *ywhaz*^-/-^ spent freezing after the addition of the second group of unfamiliar fish (p = 0.006 and p= 0.0468, respectively, Figure 7C and D) without affecting WT behaviour. However, a higher concentration of quinpirole, 1 or 4 mg/L, did not reverse the freezing behaviour (p > 0.99 for both concentrations, Figure 7D).

Several behavioural tests were repeated in a second group of adult fish and an increased freezing behaviour was observed in *ywhaz*^-/-^ fish in all the tests performed (Figure S7A). Additionally, a third batch of behavioural tests was performed in a different setup, but strong differences in the WT behaviour of this batch, probably due to environmental effects, prevented us from comparing the results obtained in this batch with the previous batches (Figure S7B).

## DISCUSSION

Functional and genetic studies had previously suggested a role for the *YWHAZ* gene in neurodevelopmental disorders such as ASD or schizophrenia (Cheah et al., 2012; Toma et al., 2014; Torrico et al., 2020; Xu et al., 2015). In this study, we have used zebrafish to investigate *ywhaz* function in neural development and behaviour, showing that *in vivo* whole-brain imaging is a powerful tool to understand the mechanisms involved in brain disorders. We found that *ywhaz* deficiency resulted in an altered hindbrain connectivity during larval stages and a neurotransmitter imbalance during adulthood leading to freezing in response to novelty.

We observed a pan-neuronal expression of *ywhaz* in larval stages, which suggests that *ywhaz* may be involved in a wide range of functions during zebrafish neurodevelopment. The important function of *Ywhaz* in neurogenesis and neuronal differentiation was previously demonstrated in mice KO models (Cheah et al., 2012; Cornell and Toyo-oka, 2017; Jaehne et al., 2015; Toyo-Oka et al., 2014; Xu et al., 2015), and disrupted neurogenesis is a known risk factor for neurodevelopmental disorders (Packer, 2016). Although expressed pan-neuronally, *ywhaz* presents a stronger expression in larvae cerebellum, a brain region whose dysfunction has been related to neurodevelopmental disorders (van der Heijden et al., 2021; Stoodley, 2016). Interestingly, decreased *YWHAZ* expression was reported in the cerebellum of ASD patients (Torrico et al., 2020). In adult zebrafish, *ywhaz* is specifically expressed in Purkinje cells (PC) in the cerebellum. *Postmortem* studies have shown a reduction in PC number, size and density in the brain of autistic patients (Bailey et al., 1998; Fatemi et al., 2002; Palmen et al., 2004; Skefos et al., 2014), and *Tsc1* mutant mice with a decrease in PC functioning show autistic-like behaviours (Tsai et al., 2012). *ywhaz* is therefore expressed in brain regions that play a crucial role in neurodevelopment and whose dysfunction is related to neurodevelopmental disorders.

Spontaneous bursting activity during development is essential for the correct assembly of neural circuits (Kirkby et al., 2013; Molnár et al., 2020). This activity has been shown to control apoptosis and future connectivity in the developing cortex of mice (Blanquie et al., 2017), and in the larval zebrafish optic tectum it is involved in the processing of sensory information (Marachlian et al., 2018). In addition, resting state magnetic resonance studies have revealed an altered spontaneous functional connectivity in patients diagnosed with ASD or schizophrenia (Van Den Heuvel and Fornito, 2014; Li et al., 2021; O’Reilly et al., 2017; Rane et al., 2015; Sheffield and Barch, 2016). Here, *ywhaz*^-/-^ larvae present a decreased burst activity and synchronization in the hindbrain, the region where *ywhaz* showed a higher expression in WT animals. These results together with the observed decreased levels of DA and 5-HT in the hindbrain of *ywhaz*^*-/-*^ adults point to long-term alterations in brain function. These findings suggest that *ywhaz* is involved in establishing brain connectivity during development and that this impaired connectivity may contribute to the subsequent neurotransmitter imbalance found in adults.

Among all 14-3-3 isoforms, *YWHAZ* plays the most important role in DA synthesis (Wang et al., 2009). *YWHAZ* is known to regulate the function of TH and TPH, rate-limiting enzymes in the biosynthesis of DA and 5-HT (Aitken, 2006). In previous work, we demonstrated that a disrupting mutation in *YWHAZ* produces a loss of affinity of the protein for TH (Torrico et al., 2020). Therefore, *ywhaz* deficiency may contribute to a decrease in TH activity and a subsequent reduction in DA levels. These results are in line with the alterations in the DA system previously reported in patients with neurodevelopmental disorders such as ASD, ADHD or schizophrenia (Kesby et al., 2018; Pavǎl, 2017; Tripp and Wickens, 2009). The increased *drd2a* and *drd2b* levels we reported in *ywhaz* mutants may be due to compensatory mechanisms to overcome DA depletion. Interestingly, schizophrenia patients have a higher density of DRD2 receptors and all antipsychotic drugs used today are DRD2 antagonists (Kestler et al., 2001; Zakzanis and Hansen, 1998). In addition, upregulation of *tph2* may be a mechanism to compensate for the depletion of 5-HT we found in the hindbrain of *ywhaz*^-/-^ adults. In line with these results, altered levels of 5-HT have been reported in ASD and schizophrenia patients (Gabriele et al., 2014; Muck-Seler et al., 2004). Finally, we analysed the expression levels of *ywhae* in adult *ywhaz*^-/-^ brains, another member of the 14-3-3 gene family that form hetorodimers with *ywhaz*. Although the differences are not significant and a further study is needed, we hypothesize that upregulation of other 14-3-3 proteins may compensate for some aspects of *ywhaz* depletion.

Finally, treatment with fluoxetine, a serotonin reuptake inhibitor, and quinpirole, a selective DRD2-like receptor agonist, was able to rescue the abnormal neophobic freezing behaviour observed in *ywhaz* mutants. Similarly, in a previous study, behavioural alterations were rescued in *Ywhaz*^-/-^ mice using the antipsychotic drug clozapine, an antagonist of DA and 5-HT receptors used as a medication for schizophrenia (Ramshaw et al., 2013). 5-HT and DA are involved in sensory processing and social cognition (Fernández et al., 2018; Jacob and Nienborg, 2018), and DA plays an important role in social reward (Rademacher et al., 2016). Given 5-HT and DA function in behaviour, we hypothesize that the alterations to 5-HT and DA signalling in mutant fish are responsible for the exaggerated response to novel stimuli present in *ywhaz*^-/-^ adults.

Several strengths and limitations of this study should be discussed. First, whole-brain imaging experiments were performed in 6 dpf larvae whereas neurotransmitter levels and behaviour were investigated in adult fish due to the impossibility of performing all these experiments at the same age. Even though this constitutes a limitation, it brings useful complementary information to analyse the effect of *ywhaz* deficiency at different levels and ages. In addition, our whole-brain imaging setup did not permit us to analyse neuronal activity in ventral telencephalic and diencephalic brain regions like the ventral thalamus and hypothalamus, where important DA and 5-HT nuclei are located. Finally, the differences observed in the behavioural phenotype between batches of experiments may be due to environmental differences (between setups) and to (epi)genetic changes between fish generations. Indeed, a previous study reported differences between *Ywhaz*^-/-^ mice with a different genetic background (Xu et al., 2015), and it is well described that environment plays an important role in the onset or phenotypical expression of psychiatric and neurodevelopmental disorders.

In conclusion, our findings highlight the important role of *YWHAZ* in neurodevelopment and shed light on the neurobiological mechanisms underlying its contribution to neurodevelopmental disorders. In addition, this work highlights the use of whole-brain imaging techniques as a promising approach for the study of neurodevelopmental disorders as it provides valuable and precise information about the mechanisms underlying the observed phenotype. Finally, pharmacological rescue of altered behaviour of *ywhaz*^-/-^ fish provide some clues for the use of specific treatments to revert the associated symptomatology in neuropsychiatric disorders such as ASD or schizophrenia.

## Supporting information

Supplementary material: Tables, Figures and Videos (titles)

## AUTHORS CONTRIBUTION

N.F-C, WHJ.N and B.C. conceived and coordinated the study, N.F-C. designed the experimental approaches for whole-brain imaging, WHJ.N designed the behavioural and pharmacological approaches and B.C. designed the genetic approaches. E.A-G. designed and conducted the whole-brain imaging and behavioural experiments and wrote the paper, E.DV. designed and conducted the CRISPR/Cas9, ISH, IHC, HPLC and behavioural experiments. AMJ.Y. contributed to the HPLC experiments. JG.O. designed the pipeline and methodology for the whole-brain imaging analyses, G.C. and E.G. conducted the whole-brain imaging recordings, E.A-G. analyzed the imaging data, F.A. contributed to the whole-brain imaging analysis and P.L-A supervised the whole-brain imaging recordings. All authors discussed and commented on the manuscript.

## ACKNOWLEDGEMENTS

GCaMP6s albino zebrafish embryos were generated by the National Institute of Genetics (Japan) and obtained from Dr. Matt Parker from the University of Portsmouth, UK. The *Tg(aldoca:gap43-Venus)* line was obtained from Masahiko Hibi from the Bioscience and Biotechnology Center of Nagoya University, Japan. *Tg(olig2:egfp)*^*vu12*^ brains were obtained from the Center for Developmental Biology, UMR 5547 CNRS, Toulouse, France. Major financial support for this research was received by BC from the Spanish ‘Ministerio de Ciencia, Innovación y Universidades’ (RTI2018-100968-B-100), the ‘Ministerio de Sanidad, Servicios Sociales e Igualdad/Plan Nacional Sobre Drogas’ (PNSD-2017I050 and PNSD-2020I042), ‘Generalitat de Catalunya/AGAUR’ (2017-SGR-738), and the European Union H2020 Program [H2020/2014-2020] under grant agreements n° 667302 (CoCA) and Eat2beNICE (728018). E.A-G was supported by the Ministerio de Economía y Competitividad (Spanish Government). G.C., E.G. and P.L-A acknowledge financial support from the Spanish Ministerio de Economía y Competitividad (MINECO) through the “Severo Ochoa” program for Centres of Excellence in R&D CEX2019-000910-S), MINECO/FEDER Ramon y Cajal program (RYC-2015-17935); Laserlab-Europe EU-H2020 GA no. 871124, Fundació Privada Cellex, Fundación Mig-Puig and from the Generalitat de Catalunya through the CERCA program. F.A. acknowledges financial support from the Spanish ‘Ministerio de Ciencia, Innovación y Universidades’ (PID2019-107738RB-I00, MICINN/FEDER) and SGR (2017SGR1255).

## DECLARATION OF INTERESTS

The authors declare no competing interests.

## STAR METHODS

### RESOURCE AVAILABILITY

#### Lead contact

Further information and requests for reagents will be fulfilled by the lead contact, Ester Antón-Galindo.

#### Materials availability

The *ywhaz* zebrafish mutant line generated in this study is available upon request. This study did not generate any other unique materials or reagents.

#### Data and code availability

- All data reported in this paper will be shared by the lead contact upon request.
- Pipelines and code used to process the whole-brain imaging datasets can be found in github:// (repository will be made public prior to publication).
- Any additional information required to reanalyse the data reported in this paper is available from the lead contact upon request.

## EXPERIMENTAL MODEL AND SUBJECT DETAILS

### Zebrafish strains, care and maintenance

Adult zebrafish and larvae (*Danio rerio*) were maintained at 28.5°C on a 14:10 light-dark cycle following standard protocols. All experimental procedures were approved by a local Animal Welfare and Ethical Review board (University of Leicester and Generalitat de Catalunya). AB wild-type (WT), *Tg*(*aldoca:gap43-Venus*), *Tg*(*olig2*:*egfp*)^vu12^, *ywhaz*^*-/-*^, albino *Tg(elavl3:GCaMP6s)* and albino *Tg(elavl3:GCaMP6s)ywhaz*^*-/-*^ zebrafish lines were used for the experiments. The *Tg(aldoca:gap43-Venus)* line was obtained from Masahiko Hibi from the Bioscience and Biotechnology Center of Nagoya University (Japan), *Tg(olig2:egfp)*^*vu12*^ brains were obtained from the Center for Developmental Biology, UMR 5547 CNRS, Toulouse (France), *ywhaz*^-/-^ line was generated by using CRISPR/Cas9 targeted mutagenesis, and albino *Tg(elavl3:GCaMP6s)* embryos were generated at the National Institute of Genetics (Japan)(Kawakami et al., 2004; Urasaki et al., 2006).

### Generation of *ywhaz* zebrafish knock out using CRISPR/Cas9

#### Design of the target sgRNA

A synthetic guide RNA (sgRNA) was designed using the online available software CHOPCHOP (Labun et al., 2016; Montague et al., 2014). We selected a target sequence containing the PAM motif within the third exon of the *ywhaz* gene and we designed the sgRNA based on this sequence: 5’-GGGTGACTATTACCGCTACC -3’ (Figure S1).

We extracted genomic DNA from 10 adult zebrafish (5 males and 5 females) and amplified the area around the CRISPR target site by PCR (forward (Fw) primer 5’-TGACCTGGTTTCTGAGCTGA -3’ and reverse (Rv) primer 5’-TGCTGAACATCAAAGACCATCT -3’). Then, we used the T7 endonuclease I (T7EI) assay to check if any mutation was present in and around the target site. 5 μl of PCR product derived from each male was mixed with 5 μl of genomic DNA from each female and the samples were diluted to a final volume of 20 μl. We denatured the samples and then reanneal them with the addition of 2 μl of NEBuffer 2 (New England Biolabs) using a thermocycler and the following protocol (95°C, 5 min; 95-85°C at −2°C/s; 85-25°C at −0.1°C/s; hold at 4°C). Hybridized PCR products were then treated with 2 U T7EI at 37°C for 1 hour in a reaction volume of 20 μl. The action of the T7EI was visualized by gel electrophoresis and the absence of SNPs in this amplified fragment was confirmed.

#### Production of sgRNA

We used the pDR274 vector, a gift from Keith Joung (Addgene plasmid #42250) (Hwang et al., 2013), to generate the template for sgRNA transcription. In order to construct a plasmid containing the designed sgRNA, the vector was digested with *BsaI* restriction enzyme (New England Biolabs), dephosphorylated using Antarctic phosphatase (New England Biolabs), and purified from a gel. Two oligonucleotides 5’-TAGGGTGACTATTACCGCTACC -3’ and 5’-AAACGGTAGCGGTAATAGTCAC-3’ were designed to generate sgRNA. Oligonucleotides were phosphorylated using T4 Polynucleotide Kinase (New England Biolabs) and annealed by incubation at 95°C for 5 minutes followed by slowly decreasing the temperature. The annealed oligonucleotides were then cloned into the vector backbone using the *BsaI* restriction site.

The sgRNA was transcribed using the mMESSAGE mMACHINE Kit (Life Technologies) and the *DraI*-digested gRNA expression vector as a template. The sgRNA was DNase treated and precipitated with ammonium acetate/ethanol following standard procedures. The RNA concentration was quantified, diluted to 100 ng/μl and stored at -80°C.

#### Production of Cas9 mRNA

*Cas9* mRNA was transcribed from the pMLM3613 vector (Addgene plasmid #42251, Keith Joung) (Hwang et al., 2013) using a mMESSAGE mMACHINE T7 Ultra Kit (Life Technologies). This vector has a unique *PmeI* restriction site positioned 3’ at the end of the Cas9 coding sequence. The *Cas9* mRNA was DNase treated and precipitated with ammonium acetate/ethanol following standard procedures. The RNA concentration was quantified, diluted to 500 ng/μl and stored at -80°C.

#### Microinjection of sgRNA/Cas9 in one-cell stage embryos

Embryos were collected by natural mating of pairs of WT zebrafish. 200-250 one cell stage embryos were co-injected with approximately 1 nl total volume of Cas9-encoding mRNA (250 ng/μl) and sgRNA (25 ng/μl) each. The day after microinjection, ten normally developed 24 hpf embryos were selected to check targeting efficiency using T7EI assay

#### Embryonic genomic DNA extraction and T7EI assay

Genomic DNA was extracted from ten 24 hpf injected embryos and every individual sample was mixed with WT genomic DNA. For genomic DNA extraction a single embryo was placed into a microcentrifuge tube containing 20 μl of base solution (25 mM NaOH and 0.2 mM EDTA) and heated to 95°Cfor 30 minutes. The tube was then cooled to 4°C and 20 μl of neutralization buffer (40mM Tris-HCl pH 5.0) were added. The sample was centrifuged and the supernatant was used directly as template for PCR. We checked the occurrence of mutations and the efficiency of the method by using the T7EI assay. Once the positive outcome of the procedure was confirmed, the rest of the injected embryos were raised to adulthood to form the F0 generation.

#### Screening by amplicon sequencing

In order to identify a founder carrying a mutation of interest that could be transmitted through the germline, we outcrossed a single F0 fish with a WT fish and collected groups of five embryos produced by each breeding pair. After the extraction of genomic DNA from these pools of embryos (40 μl of base solution were used to cover the embryos and 40 μl of neutralization buffer were subsequently added, as previously described), the samples were analysed by MiSeq Illumina sequencing. Specific primers containing a partial Illumina adaptor sequence were designed (Fw, 5’-<u>TCGTCGGCAGCGTCAGATGTGTATAAGAGACAG</u>CATCTGCTGGACAAGTTTCTGA-3’; Rv, 5’-<u>GTCTCGTGGGCTCGGAGATGTGTATAAGAGACAG</u>ATAATTGGTGTCCGGGTCAAAC-3’) and a region of about 230 bp surrounding the CRISPR target site was amplified by PCR. The quality and size of the amplicons produced by PCR were checked by running 5 μl of each PCR product on an agarose gel. In case of good quality and right size the rest of the samples were purified using a PCR & DNA Cleanup Kit (5 μg) (MonarchR, New England Biolabs). The DNA concentrations were quantified using a Qubit spectrophotometer and the amplicons were diluted to 15-25 ng/μl prior to analysis. Samples were sequenced using a MiSeq Illumina platform in collaboration with Dr Jason Rihel, University College London, UK. The read out of this analysis were FastQ data files, which were analysed using Geneious software. Once a F0 fish carrying an interesting indel transmitted to the germline was identified, the F1 embryos born from the cross between the selected F0 founder and WT were raised to adulthood. F1 fish were then tested with the T7EI assay and genotyped. Only F1 carrying the mutation were kept. We tested 40 F1 fish with the T7EI assay and found five fish carrying the 7 bp mutation (Figure S1D).

#### Genotyping

Genotyping was performed by PCR reaction. To discriminate between WT, *ywhaz*^+/-^ and *ywhaz*^-/-^ fish, we designed specific primers that recognise the presence or the absence of the Δ7 allele. We also designed a pair of control primers to confirm the efficiency of the PCR reaction (Table S1). To isolate genomic DNA for the PCR reaction from the fin, adult zebrafish were previously anesthetized in 0.02% tricaine methanesulfonate.

#### Checking for the presence of off-target cleavage sites and generation of a stable mutant line

To check the absence of any off-target cleavage before crossing F1, we used the CHOPCHOP web tool. We found two potential off-target sites for our sgRNA (Table S2), that were amplified by PCR in our selected F1 fish using the following primers: 5’-CCTCTTGTCAGTGGCGTACA -3’ (Fw) and 5’-TTCTCGGAGGACTCAACCAC -3’ (Rv), for the off-target site present in *ywhag2*, and 5’-GCAGAGGAATTACGGATGGA -3’ (Fw) and 5’-CGCGTTTATCCTGAGCTTTC -3’ (Rv) for the off-target site present in *tspan9b*. PCR products were purified using a PCR & DNA Cleanup Kit (5 μg) (MonarchR, New England Biolabs), and analysed by Sanger sequencing. No extra mutations were found in the off-target areas of the gene (data not shown) and the selected F1 fish were in-crossed to obtain a final F2 stable mutant line homozygous for the mutation of interest.

### Transgenic zebrafish lines for brain imaging experiments

*Tg(elavl3:GCaMP6s)* transgenic zebrafish larvae in the albino background (Kawakami et al., 2004; Urasaki et al., 2006) were used for brain imaging experiments as the WT group. To obtain a transgenic line expressing GCaMP6s pan-neuronally, with the albino background and not expressing *ywhaz*, we performed several crosses between the *Tg(elavl3:GCaMP6s)* and *ywhaz*^-/-^ lines. First, F0 albino *Tg(elavl3:GCaMP6s)* fish were crossed with F0 *ywhaz*^-/-^ zebrafish. F1 fish obtained from this breeding were crossed with albino *Tg(elavl3:GCaMP6s)* and F2 fish obtained from this breeding were genotyped to select only albino *Tg(elavl3:GCaMP6s) ywhaz*^+/-^ fish. Protocol used for *ywhaz* genotyping is described above. To select *Tg(elavl3:GCaMP6s)* fish we performed a specific PCR that amplifies selectively a GCaMP6s fragment and fluorescence of larvae was confirmed with a fluorescence microscope. Selected F2 fish were in-crossed and F3 fish were genotyped to select albino *Tg(elavl3:GCaMP6s) ywhaz*^-/-^ fish. F3 fish were then in-crossed to obtain a stable transgenic mutant line.

## METHOD DETAILS

### In situ hybridization (ISH)

#### Preparation of ywhaz mRNA probe

Total RNA (tRNA) was extracted from whole frozen adult WT zebrafish brains using the TRIzol reagent. Complementary DNA (cDNA) was synthesized from 1 μg tRNA using the RevertAid First Strand cDNA Synthesis Kit (Thermo Scientific). The *ywhaz* mRNA sequence was taken from the National Center for Biotechnology Information (NCBI) web site (NCBI Reference Sequence: NM_212757.2). Primers to amplify the open reading frame (ORF) of the gene were designed as follows: forward (Fw) primer 5’-AACCTGCTCTCTGTGGCCTA -3’ and reverse (Rv) primer 5’-GCTCAGAAATGGCATCATCA -3’. The 481 bp *ywhaz* amplicon was generated by PCR reaction and cloned into a plasmid using the StrataClone PCR cloning Kit (Agilent). The plasmids were collected and purified using GeneJET Plasmid Maxiprep Kit (Thermo Scientific) and the product was sequenced by GATC Biotech to check the orientation of the insert in the plasmid and the identity of the sequence. The StrataClone PCR Cloning Vector pSCA-Amp/Kan containing the *ywhaz* insert was linearized with *NotI* restriction enzyme and *ywhaz* DIG-antisense RNA probe was then generated by in vitro transcription. The product was DNase treated and cleaned using sodium acetate/ethanol precipitation. The final *ywhaz* probe was stored at -20°C.

#### Preparation of the samples for ISH

Embryos were treated with 1-phenyl 2-thiourea (PTU) at 24 hours post fertilization (hpf) to prevent pigmentation. Embryos, larvae and dissected brains from adult fish were fixed overnight at 4°C in 4% paraformaldehyde (PFA) in phosphate-buffered saline (PBS). Specimens were then dehydrated with a gradient of methanol/PBS (25%, 50%, 75% and 100% methanol) before being stored for at least one hour and up to several months at -20°C.

#### ISH protocol

##### First day of ISH

Samples were rehydrated with a gradient of methanol/PBS (75%, 50%, 25% and 0% methanol) and then digested with proteinase K (10 μg/ml in PBS) at room temperature (30 minutes for a whole adult brain and 9 dpf embryos, 25 minutes for 6 dpf embryos, 15 minutes for 3 dpf embryos and 10 minutes for 2 dpf embryos). Samples were then fixed in 4% PFA for 20 minutes and rinsed in phosphate-buffered saline + 0.1% Tween-20 (PBT). Samples were prehybridized at 68°C for at least 2 hours in 300 μl HYB+ buffer (65% formamide, 5X saline-sodium citrate (SSC) buffer, 50 μg/ml heparin, 0.5 mg/ml torula RNA, 0.1% Tween-20, 9.2 mM citric acid, pH 6.0). HYB+ was then replaced with fresh HYB+ buffer containing the DIG-labelled probe (5 ng/μl) and incubated overnight at 68°C.

##### Second day of ISH

The HYB+/probe mix was removed and stored at -20°C for future use. Samples were washed with a gradient of HYB_+_/2X SSC (75%, 50%, 25% and 0% HYB) for 10 minutes each, and then twice with 0.05X SSC for 30 minutes each. For ISH on sections, adult brains were fixed for 20 min with 4% PFA and embedded in 3% agarose dissolved in water. Samples were sectioned at 100 μm using a vibratome and sections were collected in PBS. Specimens were blocked for one hour at room temperature (RT) in blocking solution (2% normal goat serum, 2 mg/ml bovine serum albumin in PBT) and then incubated overnight with anti-DIG-AP antibody (1:4000 dilution in blocking solution).

##### Third day of ISH

Samples were washed several times in PBT and then three times for 10 minutes each in Xpho solution (100 mM Tris-HCl pH 9.5, 50 mM MgCl_2_, 100 mM NaCl and 0.1% Tween-20). Xpho solution was replaced with NBT/BCIP solution (225 μg/ml of NBT and 175μg/ml of BCIP in Xpho) and the specimens were incubated in the dark to develop the stain. Samples were monitored with a dissecting microscope every 30 minutes. The reaction was stopped by several washes in PBS and were fixed in 4% PFA for 20 minutes. Embryos were stored in 80% glycerol and 20% PBT at 4°C, whereas sections were mounted on slides and covered with Mowiol solution. The NBT/BCIP signal was imaged using a GX microscope, a CMEX 5.0 camera and Image focus 4 software.

### Immunohistochemistry (IHC)

Adult brains were rehydrated with a gradient of methanol/PBS (75%, 50%, 25%, 0% methanol) and then digested with proteinase K (10 μg/ml in PBS) at room temperature for 30 minutes followed by post-fixation in 4% PFA for 20 minutes. Samples were then rinsed in PBT. Specimens were embedded in 3% agarose dissolved in water and sectioned at 100 μm using a vibratome. Sections from *Tg*(*olig2*:*egfp*)^vu12^ were blocked for 1 hour at RT with blocking solution (1:10 normal horse serum in blocking diluent, which is 1% bovine serum albumin (BSA) in PBT) and then incubated overnight in primary antibody. The day after, samples were washed several times in PBT, blocked with blocking solution for an hour at RT and incubated in secondary antibody diluted in blocking solution for 1 hour at RT. For *ywhaz*^-/-^ and *Tg*(*aldoca*:*gap43*-*Venus*)^rk22^ the Vectastain universal *Elite* ABC Kit (Vector, #PK-6200) was used. Therefore, the sections were blocked for 1 hour at RT with the blocking serum and exposed overnight to primary antibody, followed by 1 hour incubation in secondary antibody and 3 hours’ incubation with Vectastain *Elite* ABC Reagent solution. After several washes in PBT, peroxidase activity was detected using 3,3ʹ-Diaminobenzidine (DAB, Sigma, #D4293) according to the manufacturer’s instruction. For double in situ hybridization/antibody labelling, the ISH was performed first (with NBT/BCIP staining) followed by the IHC (with DAB staining). Primary and secondary antibody concentration are specified in Table S3.

### Gene expression analysis using RT-qPCR

Total RNA was using the GenEluteTM Mammalian Total RNA Miniprep Kit (Sigma-Aldrich). Samples were DNase treated with TURBOTM DNase (ThermoFisher Scientific) to remove any genomic DNA contamination and checked for degradation. First strand cDNA was synthesised from 0.25 μg of tRNA using the RevertAid First Strand cDNA Synthesis Kit (ThermoFisher Scientific).

RT-qPCR was performed on 10 brains per genotype (WT and *ywhaz*^-/-^) with three replicates for each brain using a CFX ConnectTM Real-Time System machine (BioRad Laboratories), the SensiFASTTM SYBR No-ROX Mix (Bioline), and the following primers: Fw 5’-GAGTACCGTGAGAAGATCGAAGC -3’ and Rv 5’-CGGATCAGAAACTTGTCCAGCAG -3’ (NCBI Reference

Sequence: NM_212757). A melting curve step (50–95°C) was performed to verify that only single products had been amplified. No-template and no-reverse transcriptase controls were also performed for each primer pair and cDNA, respectively. To assess RT-qPCR efficiency, a 2-fold dilution series of cDNA template were processed. For normalization, expression levels of ribosomal protein L13a (*rpl13*) and elongation factor 1a (*elf1a*) were used as reference. The relative expression of the genes and the fold change were calculated using the 2^-ddCT^ comparative method (Livak and Schmittgen, 2001; Schmittgen and Livak, 2008).

### Whole-brain imaging

#### Light sheet microscopy recordings

Whole-brain imaging experiments were performed on 6 days-post-fertilization (dpf) albino *Tg(elavl3:GCaMP6s)* and *Tg(elavl3:GCaMP6s) ywhaz*^*-/-*^ zebrafish larvae. Larvae were first paralyzed for 10 minutes in a 1 mg/ml α-bungarotoxin solution (Thermofisher). They were subsequently placed inside a fluorinated ethylene-propylene (FEP) tube (0.7 mm inner diameter), with water and E3 medium, and then into a custom-made chamber to orientate the larvae appropriately towards the objective of the light-sheet microscope (Figure 3A).

Whole-brain imaging was performed using a custom-build Light Sheet Fluorescence Microscopy (LSFM) system in a configuration known as inverted Selective Plane Illumination Microscopy (iSPIM) (Y et al., 2011). To generate the light sheet, the laser beam from an ArKr laser (Innova 70C Spectrum, Coherent) passes through an acousto-optic tuneable filter (AOTFnC-400.650-TN, AAOptoelectronics) to select the desired wavelength (488nm) and to control the beam power. The Gaussian beam from the laser is expanded with a Galilean telescope (AC254-050-A-ML and AC254-150-A-ML, Thorlabs), and sent to the illumination arm of the iSPIM system. This is composed of a cylindrical lens (f=75mm, LJ1703RM-A, Thorlabs) forming the light sheet, a relay lens (AC254-150-A-ML and AC254-200-A-ML, Thorlabs) and the illumination objective (0.30 NA, 3.5 mm WD, Nikon CFI Plan Fluorite). Care was taken to place all the optical elements in the proper conjugated planes. A galvanometric mirror (GVS002, Thorlabs) was introduced after the cylindrical lens to scan the light sheet perpendicularly to the axis of the detection arm. The light sheet at the sample plane had a thickness of *4*.*4* µm in the entire FoV.

The detection path is composed of a 20X Objective (NA=1, WD=2mm, Olympus XLUMPLFLN), a 200 mm tube lens (TTL-200-A, Thorlabs) and a relay lens (M=0.5, AC508-200-A-ML and AC508-100-A-ML, Thorlabs). An electrically tunable lens (ETL, Optotune, EL-16-40-TC-VIS-20D) was placed at the Fourier plane of this relay lens. The ETL was used to follow the light sheet displacement created by the galvanometric mirror, keeping the generated image always in focus (Fahrbach et al., 2013). Finally, a GFP filter (Semrock, FF01-525/45-25) was placed before detection using a sCMOS camera (Hamamatsu Orca-Flash4.0 v3). The total magnification is 11.1x with and a field of view of 600µm and a pixel size of 0.5850µm.

The chamber with the Fluorinated ethylene propylene (FEP) tube (ID=0.7 mm OD=1mm) containing the zebrafish embryo was mounted into the setup, on top of a platform connected to a xyz motorized stage (NSA12, Newport). The FEP tube capillary was connected, using a screw, to a stepper motor (L4018S1204-M6, Nanotec), allowing for sample positioning and alignment with the light sheet axes. The x, y, z, position of the detection objective and the sample orientation were adjusted manually. The region of interest (ROI) was selected once the whole brain of the larva was visible having the two hemispheres displayed symmetrically (indicating a horizontal position of the larva). Once an ROI was selected, the fluorescence signal was recorded for 20 min at a scanning speed of 1 volumes/second, having 3 ms exposure time per frames, every volume had a depth of 300 µm, contained 60 imaged planes. The size of each voxel was of 0.585µm x 0.585µm x 5µm (x, y, z). See also Supplementary Videos 1-3.

#### Image segmentation and extraction of calcium signals

To extract ΔF/F traces from our calcium imaging recordings, we followed the CaIman MATLAB pipeline (Giovannucci et al., 2019). First, volumetric data were motion corrected and separated into time series for each imaged plane. Then, individual planes were processed using constrained non-negative matrix factorization method (CNMF) (Pnevmatikakis et al., 2016). We overestimated the number of ROIs, as the program discards ROIs during the refinement process, and applied a mask to avoid signal detection outside the brain. The data from each plane were then combined to produce a single dataset for an imaged larva (Figure 3B).

#### Activity and functional connectivity analysis

We used Netcal (www.itsnetcal.com) (Orlandi et al., 2017) to analyse fluorescence data and obtain single-cell and collective level statistics. ΔF/F traces were deconvolved to inferred spikes using OASIS deconvolution (Friedrich et al., 2017). To perform the analysis in different areas of the brain, we used MATLAB Volume Segmenter and the Z-brain atlas as a reference (http://engertlab.fas.harvard.edu/Z-Brain/) to delimitate five brain regions: thalamus, tegmentum, optic tectum, cerebellum and medulla oblongata (MO). We also performed analysis of network connectivity inside each of the delimited areas (Figure 3B).

### Single cell activity statistics

The number of active cells for each area was defined as the number of ROI found through the CNMF procedure mentioned above and corrected for overlapping ROI and inactive cells. ROIs that overlapped more than 20% of their volume were merged into a single ROI if the Pearson correlation between their fluorescence traces was above 0.7. Cells were deemed inactive if we could not find more than one spike after the spike deconvolution process.

A single-cell burst was defined as any set of consecutive spikes within 1 s of each other (based on the imaging framerate). The fraction of spikes within bursts was defined as the relative number of total spikes that belonged to single-cell bursts.

### Collective activity statistics

Network-wide bursts within an area were defined using a rise-detector procedure. First, the derivative of the firing rate within each neuron was computed using finite differences and averaged across the population, defining the population rise activity. A collective burst was then identified when both the average activity and the rise activity were above a given threshold (based on percentiles of each distribution). In other words, a burst was defined as a substantial increase (followed by a decrease) of activity within the population. Within a burst we defined several statistics, including the fraction of participating cells (the fraction of cells that actually fired within a burst) and the burst amplitude (the average increase in firing rate during a burst).

Variance-explained curves were computed by performing PCA decomposition using the activity of all the neurons within a given area, and then plotting the total fraction of variance explained of the original data by sorting and adding cumulatively each of the obtained components.

### Functional connectivity analysis

Functional connectivity measures were computed by first computing the Pearson correlation coefficient between the time-series of every neuron pair within a larvae. Neuron pairs were deemed as functionally connected if their average correlation was above a given threshold (the 99^th^ percentile of the whole distribution of correlation values). Network measures on the functional connectivity matrices were computed using the Brain Connectivity Toolbox (BCT) (Rubinov and Sporns, 2010).

### High performance liquid chromatography (HPLC) analysis of monoamines and metabolites

Fish were decapitated and their brains were dissected and divided into four areas: telencephalon (Tel), diencephalon (DI), optic tectum (TeO) and hindbrain (Hb). HPLC analysis for dopamine (DA), serotonin (5-HT), 3,4-dihydroxyphenylacetic acid (DOPAC), and 5-hydroxyindoleacetic acid (5-HIAA) was carried out using HPLC with electrochemical detection (Young, 2004).

Samples were weighed, homogenised in 100 μl ice-cold 0.1 N perchloric acid using a pellet pestle (Sigma, #Z359971) and centrifuged at 12,000 rcf for 15 min at 4°C. The supernatant was collected and stored at -80°C until use.

Samples (15 μl) were then injected for HPLC analysis using a Spark Triathlon refrigerated autosampler and separation was achieved on a Luna C18(2), 5 μm, 100 Å, 100 x 1 mm (Phenomenex Ltd) reverse phase column. The mobile phase (75 mM sodium dihydrogen phosphate, 1 mM EDTA, 0.6 mM octane sulphonic acid (OSA) in deionised water containing 5% methanol, pH 3.7) was delivered by a Rheos 4000 pump (Presearch, UK). Electrochemical detection was performed using a glass carbon working electrode set at 700 mV relative to an Ag/AgCl reference electrode, using an Antec Intro detector incorporating a low volume (VT-03) flow cell (Antec, Netherlands).

Samples were quantified by comparison with standard solutions of known concentrations of monoamines and metabolites using Chrom Perfect data analysis software (Justice Laboratories, NJ). Each sample was run in duplicate and the mean content of monoamine and neurotransmitter for sample was calculated and normalised to the weight of tissue. Results are expressed as picomoles per milligram of brain tissue.

### Behavioural tests

A battery of behavioural tests was performed in adult zebrafish (3-5 months-old) mixed groups of both sexes. Two tests to assess social behaviour were also performed on juvenile (one month-old) zebrafish. All fish were genotyped, sized-matched and maintained in groups by genotype until the day of testing. All behavioural experiments were completed between 10:00 and 17:00 and recorded using FlyCapture2 2.5.2.3 software and a digital camera from Point Grey Research. Fish were left for 30 minutes to habituate to the testing room before the experiment. Adult fish were tested for social behaviour, anxiety, locomotion and aggression, and juvenile fish were tested for social behaviour. For adult fish: n = 12/genotype (1^st^ batch), n=14 WT and 13 *ywhaz*^-/-^ (2^nd^ batch) and n = 13/genotype (3^rd^ batch). For juvenile fish: n = 10/genotype.

#### Visually-mediated social preference test (VMSP)

The experiment was performed in two steps as described in Carreño Gutierrez et al., 2019 (Carreño Gutiérrez et al., 2019). We used a mixture of size-matched males and females as strangers since they can attract both male and female zebrafish (Ruhl et al., 2009). Time spent in different zones of the tank was quantified using Noldus Ethovision XT software. The same procedure was used to test juvenile fish using a similar tank with a 1:3 reduction in size. The effect of novel non-social stimuli was also tested using marbles instead of the second group of unfamiliar fish in the second step of the test.

#### Shoaling test

The shoaling experiment was performed following the protocol from Parker et al., 2013 (Parker et al., 2013). We used ViewPoint Life Sciences to measure the nearest neighbour distance (NDD) and the inter-individual distance (IID). We also virtually divided the tank into eight equal sections and calculated the cluster score (Parker et al., 2013). The same procedure was used to test juvenile fish using a similar tank with a 1:3 reduction in size. Adult fish: n = 2 groups of 5 fish/genotype. Juvenile fish: n = 4 groups of 5 fish/genotype.

#### Novel tank test (NTT)

The NTT was performed in a standard 1.5 L trapezoid tank (Egan et al., 2009) and fish were recorded for 5 minutes. Noldus Ethovision XT software was used to measure the amount of time spent in the bottom (geotaxis) and top third of the tank, the time spent freezing, the total distance swum, the velocity and the absolute angular velocity.

#### Open field test

The open field test was performed in a large open tank (20.5 cm x 29 cm x 19 cm) and fish were recorded from above for 5 minutes. We used Noldus Ethovision XT software to quantify the duration of thigmotaxis (time spent swimming at a distance of 2 cm or less from the walls), the time spent in the periphery and in the centre, the time spent freezing, the distance swum and the velocity.

#### Aggression test

Aggression was measured using the mirror-induced stimulation protocol (Gerlai et al., 2000; Norton et al., 2011). Fish were placed in holding tanks with walls covered in white paper on the night before the experiment to habituate them to the setup. Single fish were recorded for 5 minutes. The time spent in agonistic interaction was manually quantified using LabWatcher software from ViewPoint Life Sciences.

#### Black and white test

The black and white test was performed in a rectangular tank (24 cm x 12 cm) divided into two equal areas, a black area and a white area. Fish were placed in the centre of the tank and recorded for 5 minutes. The time spent in each area was manually quantified.

### Drug treatments

Fluoxetine 5 mg/L diluted in DMSO (Tocris #0927) was administered by immersion for 2 hours. Quinpirole 0.25-4 mg/L diluted in H_2_O (Tocris #1061) was administered by immersion for 1 hour.

## QUANTIFICATION AND STATISTICAL ANALYSIS

Statistical analysis of RT-qPCR, HPLC and behavioural data were performed in GraphPad Prism 6. The data sets were assessed for normality using D’Agostino-Pearson and Shaphiro-Wilk normality test. Either a Mann-Whitney or an unpaired t-test with Welch’s correction was performed to compare two groups, with corrections for multiple comparisons when needed.

Statistical analysis of neuronal activity and functional connectivity consisted of unpaired t-test between genotypes for each area. Individual results for each animal are always presented in the figures together with mean, 95% confidence intervals and standard deviation of the samples.

## Notes

### Competing Interest Statement

The authors have declared no competing interest.

### Summary of Updates

Material and Methods and Results sections have been updated to be more understandable. All the figures have been revised and reordered. Supplementary files updated.

