## Supplementary material: Tables, Figures and Videos (titles) for "Deficiency of the *ywhaz* gene, involved in neurodevelopmental disorders, alters brain activity and behaviour in zebrafish"

1. **SUPPLEMENTARY TABLES**
2. **SUPPLEMENTARY FIGURES**
3. **SUPPLEMENTARY VIDEOS (titles)**

### 1. SUPPLEMENTARY TABLES

**Table S1.** *ywhaz* genotyping primers designed to discriminate between WT, *ywhaz*$^{+/-}$ and *ywhaz*$^{-/-}$ fish.

| | | |
|---|---|---|
| **Specific primers for the Δ7 allele** | *ywhaz* forward (5'3') | TGACCTGGTTTCTGAGCTGA |
| | Mutant reverse (5'-3') | TGTAGCGACTTCTAGCGGT |
| **Specific primers for WT allele** | *ywhaz* forward (5'3') | TGACCTGGTTTCTGAGCTGA |
| | WT reverse (5'-3') | TAGCGACTTCTGCCAGGTAG |
| **Internal control primers** | Control forward (5'-3') | TGTACAAGTGCAGAAACCCAC |
| | Control reverse (5'-3') | TATCCGAATCAAGGCCAGGA |

**Table S2.** Off-target sequences determined by CHOPCHOP.

| **Number of mismatches** | **Off-target sequences** | **Alignment between CRISPR target sequence and off-target sequences** | **Position** | **Gene name** |
|---|---|---|---|---|
| 2 | GGGAGATTATTACCGCTACCTGG | 1 GGGTGACTATTACCGCTACCTGG 23<br>1 GGGAGATTATTACCGCTACCTGG 23 | Exonic | *ywhag2* |
| 4 | GGGGGCCTATTACGGCTCCCGGG | 1 GGGTGACTATTACCGCTACCTGG 23<br>1 GGGGGCCTATTACGGCTCCCGGG 23 | Intronic | *tspan9b* |

**Table S3.** Primary and secondary antibody concentration used for the IHC assays.

| **Primary antibody** | **Secondary antibody** | **Genotype** |
|---|---|---|
| **Anti-GFP (Amsbio, #TP401) dilution 1:500** | Peroxidase Anti-rabbit igG (H+L) (Vector, #PI-1000), dilution 1:1000 | *Tg*(*olig2*:*egfp*)$^{vu12}$ |
| **Anti-parvalbumin (Millipore, #MAB1572) dilution 1:1000** | Biotinylated Universal antibody anti-mouse and rabbit IgG (H+L) (Vectastain universal *Elite* ABC Kit, Vector) | *ywhaz*$^{-/-}$ |
| **Anti-GFP (Amsbio, #TP401) dilution 1:500** | Biotinylated Universal antibody anti-mouse and rabbit IgG (H+L) (Vectastain universal *Elite* ABC Kit, Vector) | *Tg*(*aldoca*:*gap43-Venus*)$^{rk22}$ |

### 2. SUPPLEMENTARY FIGURES

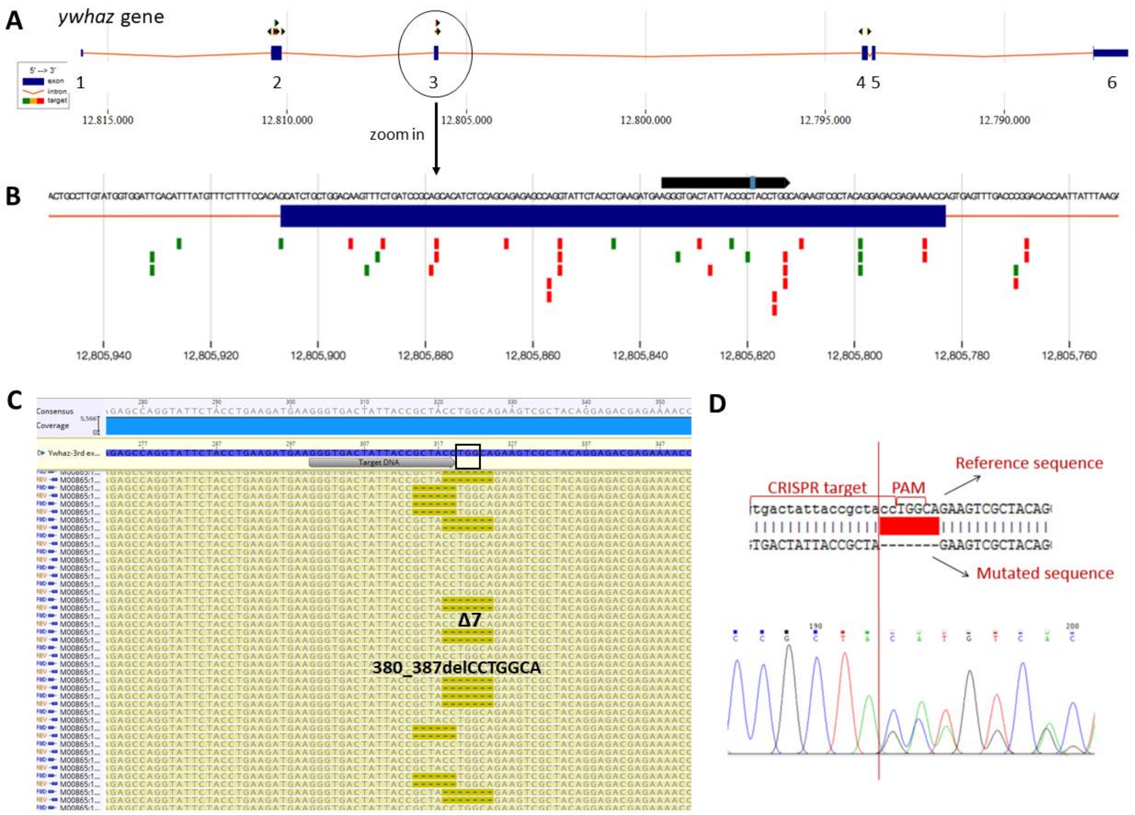


**Figure S1. Design of target sequence and sgRNA for *ywhaz* using the CHOPCHOP web tool and selected CRISPR/Cas9-mediated 7-bp deletion in *ywhaz.* Related to STAR Methods. (A)** All the possible target sequences in *ywhaz* (arrowhead) found by CHOPCHOP are shown above every exon (blue bars). Introns are represented as red lines in between exons. **(B)** The target sequence selected (black bar) within the third exon (blue bar) is represented here in greater detail. Figures are taken and adapted from http://chopchop.cbu.uib.no. **(C)** Readout of the MiSeq Illumina analyisis indicating the deletions of 6 and 7 bp in *ywhaz* caused by the microinjection of Cas9 mRNA and sgRNA. The grey arrow represents the *ywhaz* target DNA sequence, and the black rectangle the PAM sequence. MiSeq reads (yellow) are paired and aligned to the gene-specific reference sequence (blue). Dashed lines represent deletions. **(D)** Top, reference sequence containing the CRISPR target sequence and PAM motif within the third exon of *ywhaz* aligned with the correspondent sequence carrying the 7-bp deletion, indicated in red. Bottom, Sanger sequencing showing the frameshift introduced by the 7-bp deletion.

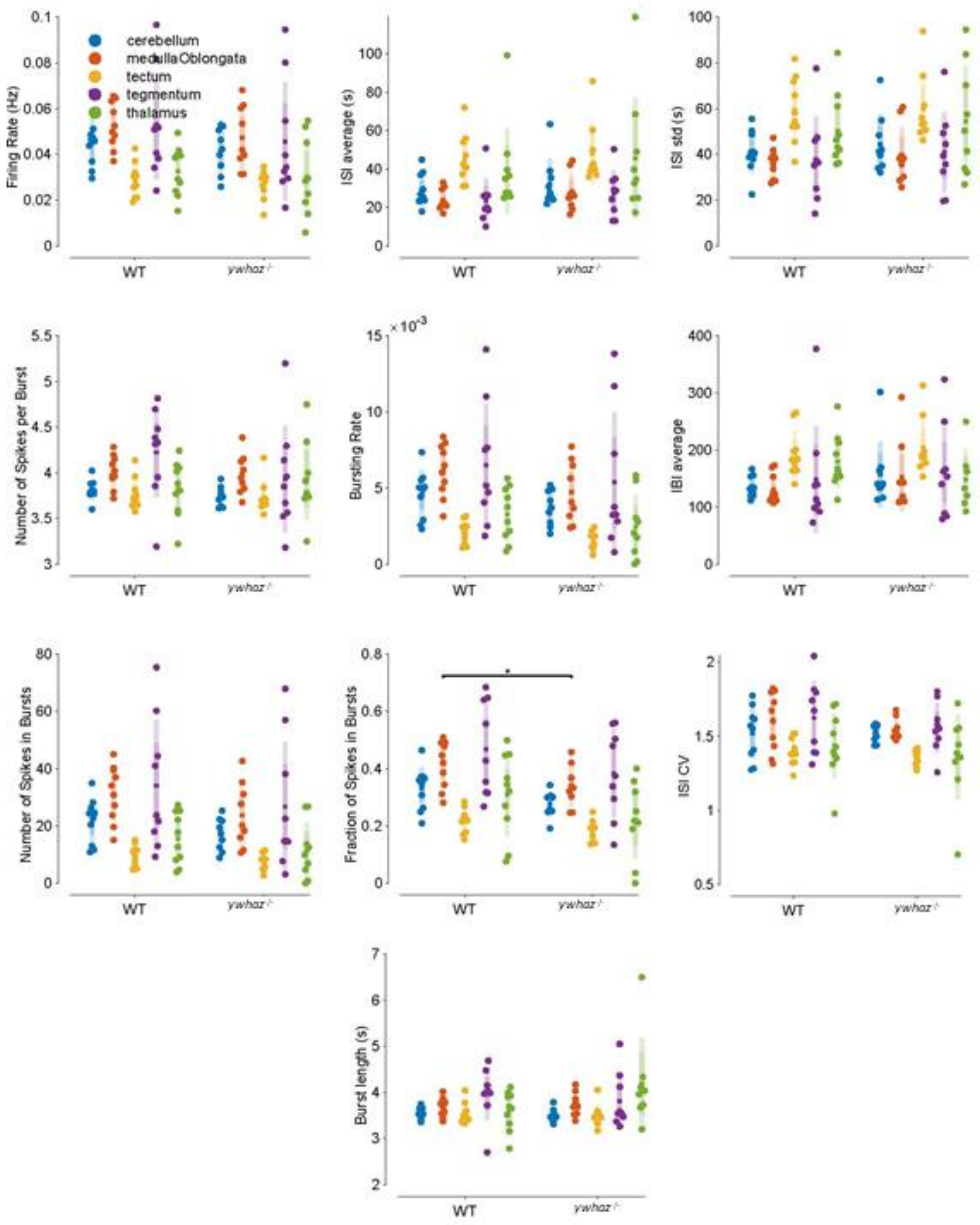


**Figure S2. Single-cell activity analysis of whole-brain imaging recordings performed in the five defined brain areas. Related to Figure 4.** n = 10 WT and 9 *ywhaz*$^{-/-}$. Unpaired t-tests, each single point represents an individual, the central line represents the mean, the darker bar the 95% confidence interval and the lighter bar the standard deviation. YWHAZ, *ywhaz*$^{-/-}$ larvae; WT, wild-type larvae. * $p < 0.05$.

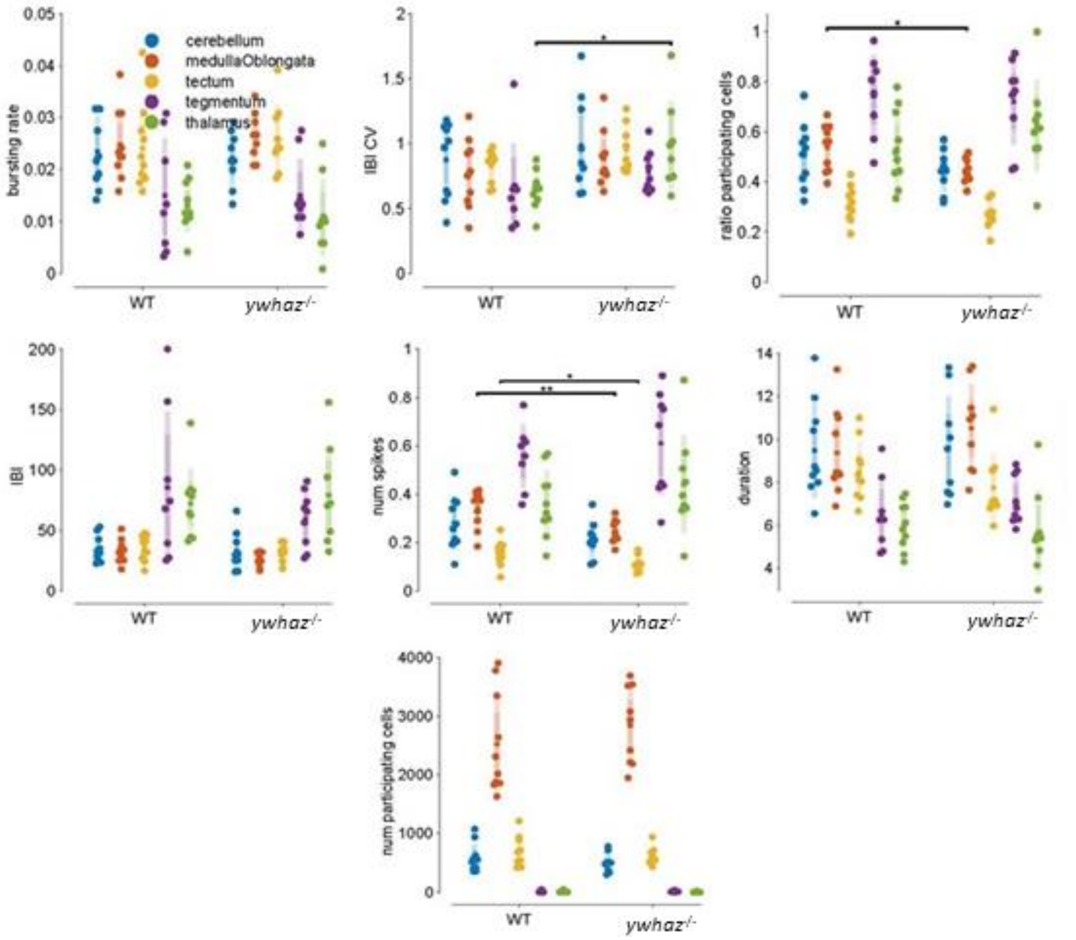


**Figure S3. Collective burst activity analysis of whole-brain imaging recordings performed in the five defined brain areas. Related to Figure 4.** n = 10 WT and 9 *ywhaz*$^{-/-}$. Unpaired t-tests, each single point represents an individual, the central line represents the mean, the darker bar the 95% confidence interval and the lighter bar the standard deviation. * $p < 0.05$, ** $p < 0.01$.

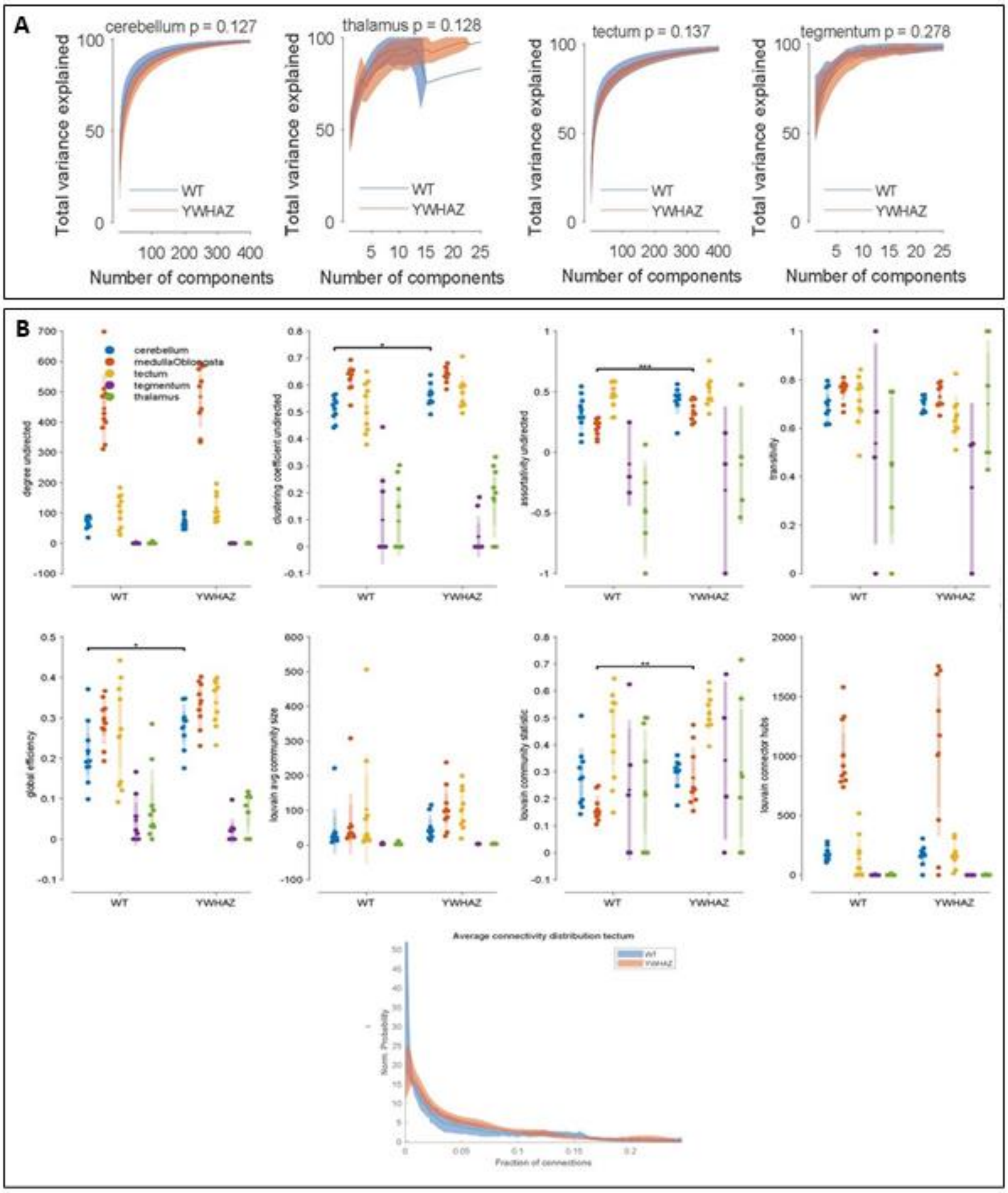


**Figure S4. Principal component and connectivity analysis of whole-brain imaging recordings. Related to Figure 5. (A)** Principal component analysis of neuronal activity performed in cerebellum, thalamus, tectum and tegmentum. YWHAZ, *ywhaz*$^{-/-}$ larvae; WT, wild-type larvae. n= 10 WT and 9 *ywhaz*$^{-/-}$. The central line represents the mean and the coloured shadow represents the 95% confidence interval. YWHAZ, *ywhaz*$^{-/-}$ larvae; WT, wild-type larvae. **(B)** Connectivity analysis performed in the five defined brain areas and neuronal connectivity distribution in the optic tectum. Top, unpaired t-tests of connectivity parameters, each single point represents an individual, the central line represents the mean, the darker bar the 95% confidence interval and the lighter bar the standard deviation. Bottom, connectivity distribution in the tectum of WT and *ywhaz*$^{-/-}$ larvae. The central line represents the mean and the coloured shadow represents the 95% confidence interval. YWHAZ, *ywhaz*$^{-/-}$ larvae; WT, wild-type larvae. n = 10 WT and 9 *ywhaz*$^{-/-}$. * $p < 0.05$, ** $p < 0.01$, *** $p < 0.001$.

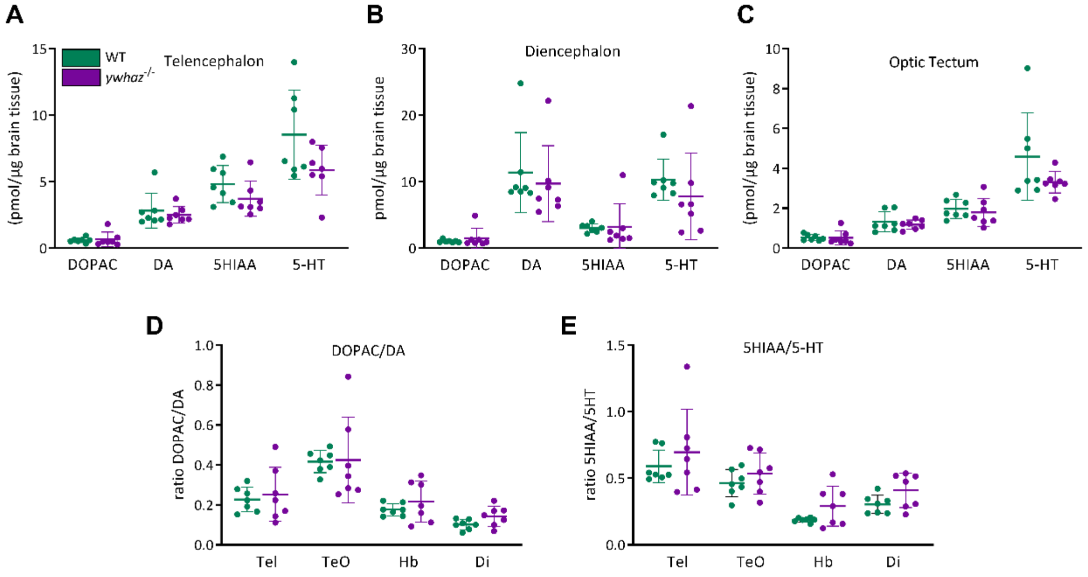


**Figure S5. Analysis of the effect of *ywhaz* loss-of-function on monoamines and their metabolites levels in different areas of WT and *ywhaz*$^{-/-}$ adult brains by high performance liquid chromatography (HPLC). Related to Figure 5.** No significant difference of monoamine levels was found in **(A)** telencephalon, **(B)** diencephalon and **(C)** optic tectum. There was also no significant difference in the breakdown of **(D)** DOPAC/DA and **(E)** 5HIAA/5-HT in any of the brain areas. Multiple t-tests with Holm-Sidak correction for multiple comparisons. Abbreviations: DA, dopamine; Di, diencephalon; DOPAC, 3,4-dihydroxyphenylacetic acid; Hb, hindbrain; Tel, telencephalon; TeO, optic tectum; 5-HIAA, 5-hydroxyindoleacetic acid; 5-HT, 5-hydroxytryptamine. n = 7 WT, n = 7 *ywhaz*-/-. Mean ± SEM.

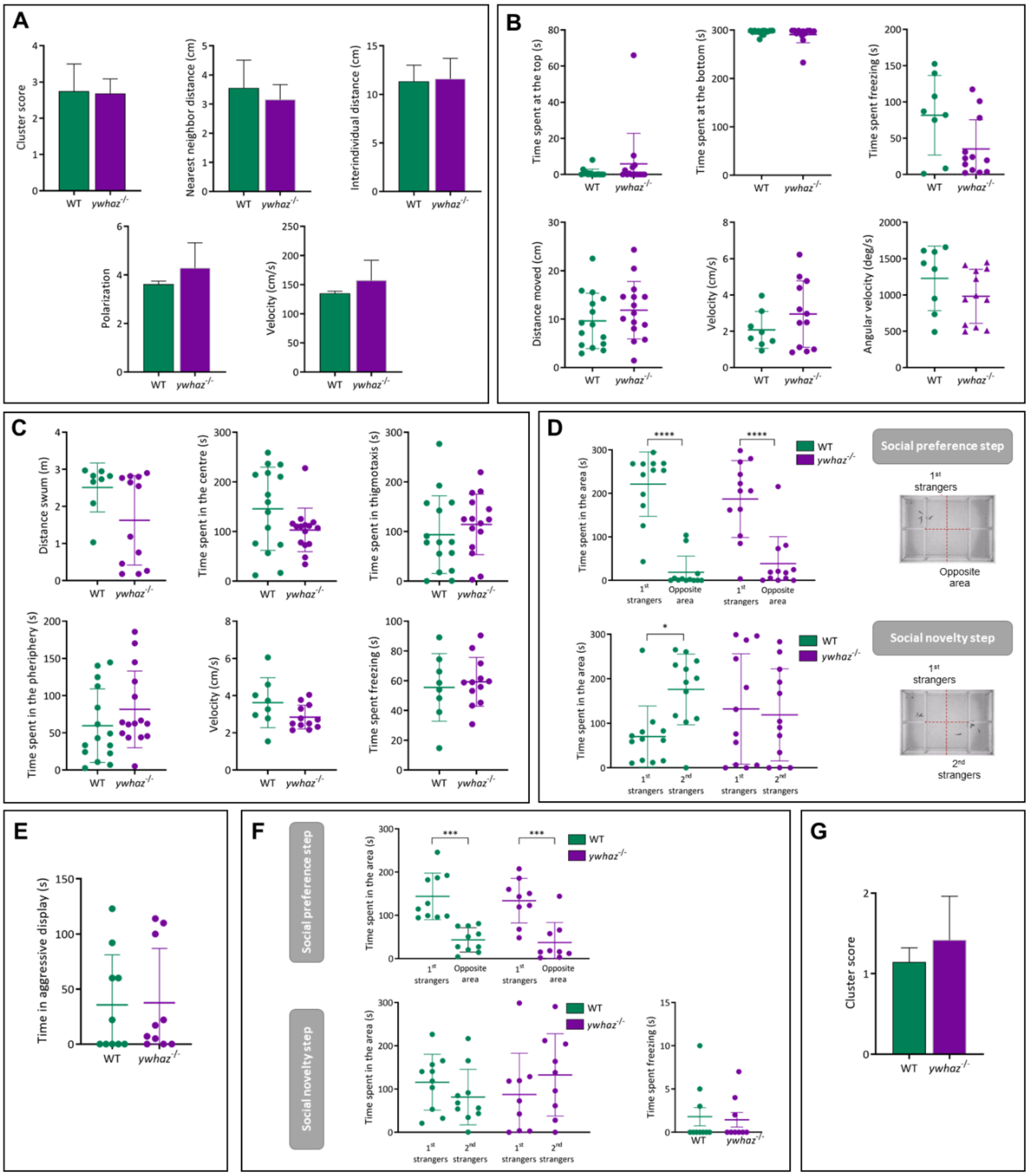


**Figure S6. Behavioural tests performed in adult (A-E) and juvenile (F-G) fish. (A) Shoaling behaviour. Related to Figure 5.** Adult *ywhaz*$^{-/-}$ display normal shoaling. Nearest neighbour distance ($p = 0.66$), inter-individual distance ($p = 0.91$), cluster score ($p = 0.93$), polarization ($p = 0.53$) and velocity ($p = 0.53$). n = 2 groups of 5 wild-type (WT), n = 2 groups of 5 *ywhaz*$^{-/-}$. **(B) Novel tank test.** Adult *ywhaz-/-* exhibit normal anxiety-like behaviour. Time spent at the bottom ($p = 0.41$), at the top ($p = 0.50$), and freezing ($p = 0.13$) in a novel tank. n = 15 WT, n = 15 *ywhaz*$^{-/-}$. Mann-Whitney U test. Locomotion ($p = 0.31$), velocity ($p = 0.19$) and angular velocity ($p = 0.22$) in a novel tank. n = 15 WT, n = 15 *ywhaz*$^{-/-}$. Unpaired t-test with Welch's correction. **(C) Open field test.** Adult *ywhaz*$^{-/-}$ behave similarly to WT in the open field test. Time at the side of the tank ($p = 0.42$), in the periphery ($p = 0.24$), in the centre of the tank ($p = 0.19$) and time spent freezing ($p = 0.70$). n = 15 WT, n = 15 *ywhaz*$^{-/-}$. Unpaired t-test with Welch's correction. Locomotion ($p = 0.057$). n = 15 WT, n = 15 *ywhaz*$^{-/-}$. Mann-Whitney U test. Velocity ($p = 0.16$) is not affected in the open field test. n = 15 WT, n = 15 *ywhaz*$^{-/-}$. Unpaired t-test with Welch's correction. **(D) Visually-mediated social preference test in adults.** On the top, social preference step: both WT and *ywhaz*$^{-/-}$ show a significant preference to spend time near a group of unfamiliar fish (1st strangers;

p < 0.0001 for both WT and *ywhaz*$^{-/-}$, two-way ANOVA with Tukey's post hoc comparisons, n = 12). On the bottom, preference for social novelty step: WT switch preference and spend more time close to the second group of unfamiliar fish (2nd strangers; p = 0.048). *ywhaz*$^{-/-}$ spend an equal amount of time near both groups of unfamiliar fish (1st and 2nd strangers; p = 0.98). Two-way ANOVA with Tukey's post hoc comparisons, n = 12. **(E) Mirror-induced aggression test**. No difference in aggression levels between adult WT and *ywhaz*$^{-/-}$ (p = 0.72). n = 15 WT, n = 15 *ywhaz*$^{-/-}$. Mann-Whitney unpaired t-test. Mean ± SEM. **(F) Visually-mediated social preference test in juvenile fish.** Top, social preference step: similar to adults, both WT and *ywhaz*$^{-/-}$ juvenile fish show a significant preference to spend time near a group of unfamiliar fish (1$^{st}$ strangers; p < 0.001 for both WT and *ywhaz*$^{-/-}$). Bottom, preference for social novelty step: WT and *ywhaz*$^{-/-}$ juveniles spend equal time close both groups of unfamiliar fish (2$^{nd}$ strangers; p = 0.077 and p = 0.63 respectively). Two-way ANOVA with Tukey's post hoc comparisons. n = 10. Time spent freezing during the first two minutes after the addition of the second group of unfamiliar fish in the behavioural tank: Neither WT nor *ywhaz*$^{-/-}$ juveniles show any freezing reaction (p > 0.99). Mann-Whitney U test. n = 10. Mean ± SEM. **(G) Cluster score calculated for juvenile fish.** Shoaling behaviour is normal in juvenile fish, with equal cluster score between genotypes (p = 0.33). n = 4 groups of 5 WT, n = 4 groups of 5 *ywhaz*$^{-/-}$. Unpaired t-tests with Welch's correction. *** p < 0.001. Mean ± SEM.

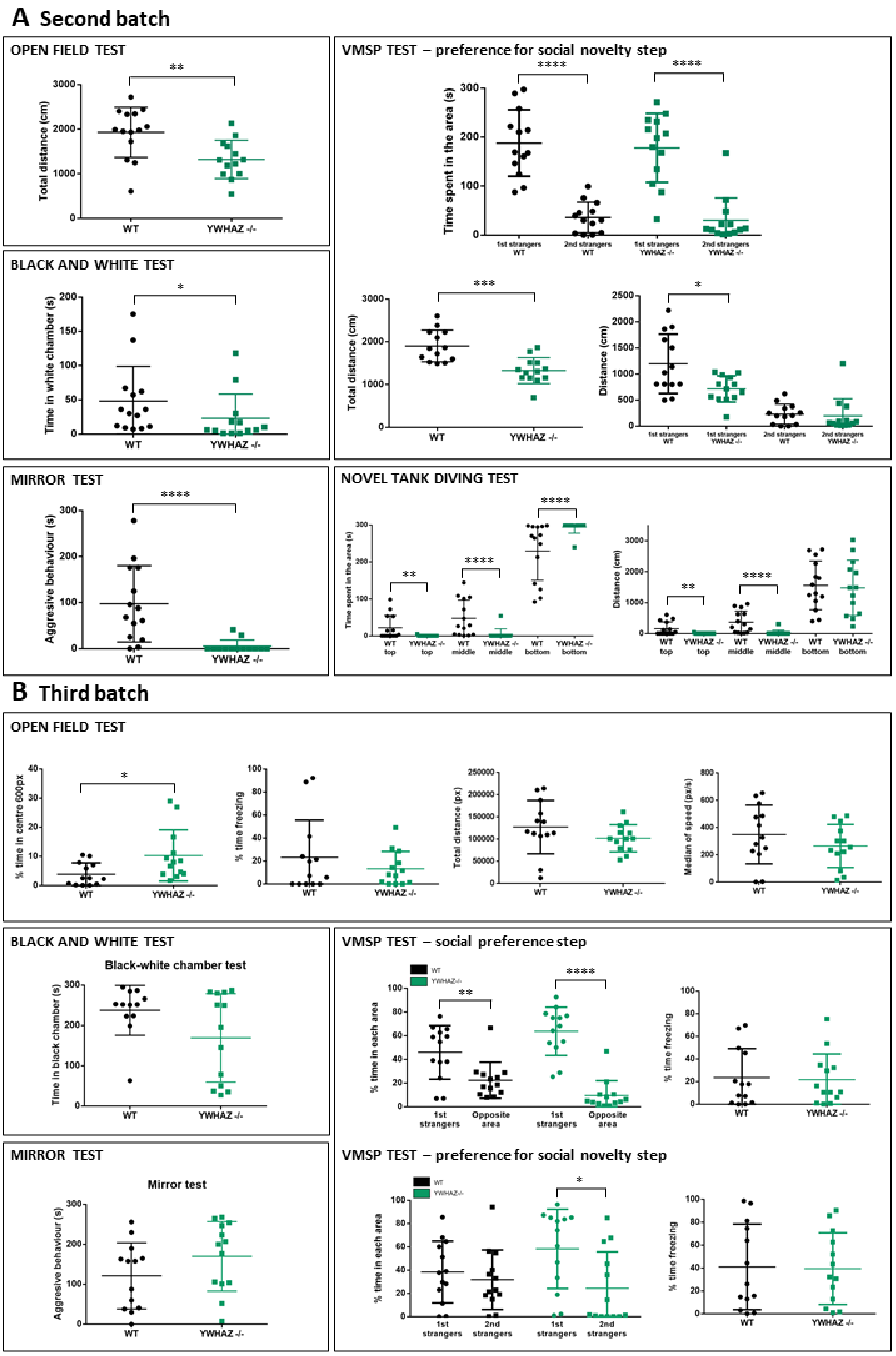


**Figure S7. Behavioural tests repeated in two other batch of adult fish. (A) Results for a second batch of adult fish. Related to Figure 5. Open field test.** Total distance travelled is significantly lower in *ywhaz*$^{-/-}$ compared to WT (p = 0.0040, Unpaired t-test with Welch's correction) due to an increased freezing observed in mutant fish **Visually-mediated social**

**preference test.** *ywhaz*$^{-/-}$ behave similarly to WT in the preference for social novelty step and show preference for the first group of strangers (Time spent in the area: 1$^{st}$ strangers vs 2$^{nd}$ strangers; p < 0.0001; *ywhaz* $^{-/-}$ p < 0.0001; Two way ANOVA, no RM, followed by Sidak's post hoc test). However, *ywhaz*$^{-/-}$ present a freezing behaviour, reflected in the lower distance travelled by mutant fish (Total distance, p = 0.0002; Distance travelled in the 1$^{st}$ strangers area, p = 0.0126). Unpaired t-tests with Welch's correction. **Black and white test.** *ywhaz*-/- spend less time in the white chamber than WT fish (p= 0.0199, Mann-Whitney U test). Indeed, they spend more time freezing in the black chamber. **Mirror-induced aggression test**. *ywhaz*$^{-/-}$ spend less time performing an aggressive behaviour (p < 0.0001, Mann-Whitney U test), as they spend most of the time freezing. **Novel tank diving test.** *ywhaz*$^{-/-}$ exhibit a higher anxiety-like behaviour: they spend more time than WT fish at the bottom area (p < 0.0001), and less time at the middle (p < 0.0001) and at the top (p = 0.0.63) areas of the novel tank. *ywhaz*$^{-/-}$ travel less distance at the middle (p < 0.0001) and top (p = 0.0050) areas of the novel tank. Mann-Whitney U tests. For all the experiments, n= 14 WT, n = 13 *ywhaz*$^{-/-}$. Mean ± SD. * p < 0.05; ** p < 0.01; *** p < 0.001; **** p<0.0001. **(B) Results from a third batch of adult fish in a different setup. Open field test.** *ywhaz*$^{-/-}$ behave similarly to WT in the open field test although they spend more time in the centre of the arena (p=0.016, Mann-Whitney U test). Time freezing (p = 0.83, Mann-Whitney U test), total distance travelled (p = 0.19, Unpaired t-test with Welch's correction) and median of speed (p = 0.27, Unpaired t-test with Welch's correction). **Black and white test.** Although no significant statistical differences (p = 0.12, Mann-Whitney U test), WT fish seem to be more anxious, as they spend almost the whole time in the black area while *ywhaz*-/- fish show no preference for any area. **Visually-mediated social preference test.** On the top, social preference step: both WT and *ywhaz*$^{-/-}$ show a significant preference to spend time near a group of unfamiliar fish (1$^{st}$ strangers vs opposite area: WT p = 0.0035, *ywhaz*$^{-/-}$ p < 0.0001; Two way ANOVA, no RM, followed by Sidak's post hoc test) and freeze a similar amount of time (p = 0.99, Mann-Whitney U test). On the bottom, preference for social novelty step: WT fish spend an equal amount of time near both groups of unfamiliar fish and ywhaz$^{-/-}$ spend more time close to the first group of strangers (1$^{st}$ strangers vs 2$^{nd}$ strangers: WT p = 0.814, *ywhaz*$^{-/-}$ p = 0.011; Two way ANOVA, no RM, followed by Sidak's post hoc test). **Mirror-induced aggression test**. No difference in aggression levels between WT and *ywhaz*$^{-/-}$ (p = 0.16, Unpaired t-test with Welch's correction). For all the experiments, n= 13 WT, n = 13 *ywhaz*$^{-/-}$. Mean ± SD. * p < 0.05; ** p < 0.01; **** p<0.0001.

### 3. SUPPLEMENTARY VIDEOS

**Supplementary video 1**. Planes of the brain covered during the *in vivo* whole-brain imaging recordings.

**Supplementary video 2.** Maximum projection of the neuronal activity during the *in vivo* whole-brain imaging recording of a single individual (15 minutes). Top, lateral and front view.

**Supplementary video 3.** 3D rotation of the maximum projection of the neuronal activity during the *in vivo* whole-brain imaging recording of a single individual.
